# Macrophage adaptation to hypoxia in the tuberculous granuloma potentiates mycobacterium-induced mitochondrial damage and granuloma necrosis

**DOI:** 10.64898/2026.02.06.702658

**Authors:** Antonio J. Pagán, Bingnan Lyu, Michal Eisenberg-Bord, Elisabeth M. Busch-Nentwich, Lalita Ramakrishnan

## Abstract

Granulomas, organized macrophage-rich aggregates, are pathological hallmarks of tuberculosis. Characteristically, as Mycobacterium tuberculosis replicates within macrophages, the granuloma enlarges, and then the center undergoes necrosis, promoting mycobacterial growth in the debris (caseum). This sequence results in large, mycobacterium-rich granulomas that can rupture into airways, facilitating disease transmission. Here, using the zebrafish-Mycobacterium marinum tuberculosis model, we find that both stages – granuloma enlargement and necrosis – depend on the mycobacterial virulence factor, EsxA, that induces mitochondrial damage and apoptosis of infected macrophages. Initially, dying macrophages are engulfed by newly recruited macrophages. Then, the enlarging granuloma core becomes hypoxic and induces Hypoxia Inducible Factor-1 (HIF-1). HIF-1 suppresses mitochondrial respiration, sensitizing macrophages to EsxA mitotoxicity. These sensitized macrophages undergo accelerated apoptosis which outstrips clearance, causing mycobacterium-rich necrotic debris to accumulate in the granuloma cores. Thus, tuberculosis transmission depends on the dynamic interplay between a mycobacterial virulence factor and an adaptive host metabolic program.

## INTRODUCTION

*Mycobacterium tuberculosis* (Mtb) and other pathogenic mycobacteria infect macrophages and induce their aggregation into granulomas, organized immunopathological structures.^1–3^ The cores of larger granulomas undergo necrotic breakdown, a critical pathogenic event because it releases mycobacteria into the cellular debris, where their growth is enhanced.^1,2,4–9^ Moreover, breakdown of the granuloma cores is more likely to result in their rupture into the airways, facilitating Mtb transmission to new hosts.^2,9,10^

Zebrafish infected with *Mycobacterium marinum* (Mm) enable the study of aspects of tuberculosis (TB) pathogenesis that are difficult or impossible to investigate in other animal models. Mm, a close relative of Mtb, causes a wasting TB-like disease in zebrafish that features organized granulomas with necrotic cores that resemble human TB granulomas.^9,11–16^ As in human TB granulomas, bacteria are abundant in the necrotic cores but sparse in the cellular regions.^14,16,17^ The optical transparency of larval zebrafish has enabled the real-time visualization of early mycobacterium-macrophage interactions leading to granuloma formation and enlargement.^3,11,18–22^ It was found that the mycobacterial virulence factor EsxA (formerly ESAT-6), a substrate of the specialized ESX-1 secretion system, promotes granuloma formation and enlargement. EsxA coordinately increases death of infected macrophages and recruitment of uninfected macrophages to the dying cells.^3,23^ EsxA-mediated death induces morphological features of apoptosis – phosphatidylserine exposure on the plasma membrane, nuclear collapse and DNA fragmentation, and cytoplasmic blebbing.^3,19^ Moreover, in keeping with apoptotic cell clearance (efferocytosis) mechanisms, the dying macrophages are phagocytosed by recruited macrophages.^24^ Cycles of infected macrophage death coupled to their efferocytosis provide mycobacteria with expanding growth niches in the macrophages that engulf them together with the dying macrophages.^3^ While this process serves to expand mycobacteria, they remain intracellular, spreading within the macrophages of enlarging granulomas.

Zebrafish granuloma breakdown, like granuloma enlargement, is EsxA-dependent.^2,14,19^ Moreover, human and mouse granulomas induced by the vaccine strain BCG, which lacks EsxA, are predominantly non-necrotic.^25–28^ Here, we have tackled the question of how EsxA both builds and breaks down the granuloma. We show that both granuloma enlargement and breakdown are due to EsxA-mediated mitotoxicity and apoptosis, which are accelerated in response to HIF-1 stabilization in the enlarging granulomas’ hypoxic cores.

## RESULTS

### EsxA-mediated mitochondrial damage and apoptotic death promote mycobacterial expansion in cellular granulomas

ESX-1 is reported to cause mitochondrial damage of cultured Mtb-infected macrophages. ^29,30^ To visualize ESX-1-mediated mitochondrial damage in granuloma macrophages in real time, we engineered transgenic zebrafish with macrophage mitochondria expressing green fluorescence (See methods). Crossing these fish with a transgenic line labeling macrophage membranes red fluorescent enabled simultaneous visualization of mitochondrial morphology and macrophage viability. Examining individual infected macrophages two days post infection (2 dpi), we could visualize their mitochondria and capture by time-lapse video microscopy their progression from an elongated to a spherical morphology, indicative of mitochondrial damage (Figure 1A). This was rapidly followed by death as evidenced by loss of plasma membrane integrity (loss of red fluorescence and dispersion of the green mitochondrial signal) (Figure 1A). Next, we assessed macrophage mitochondrial morphology in early WT granulomas. The mitochondria of the granuloma macrophages, most of which were infected as judged by colocalization with bacteria, were more spherical than those of nearby uninfected macrophages (Figure 1B, left). To determine if mitochondrial damage in granuloma macrophages was mediated by ESX-1, we assessed mitochondrial sphericity in WT and *ΔESX-1* Mm infections.

**Figure 1.**
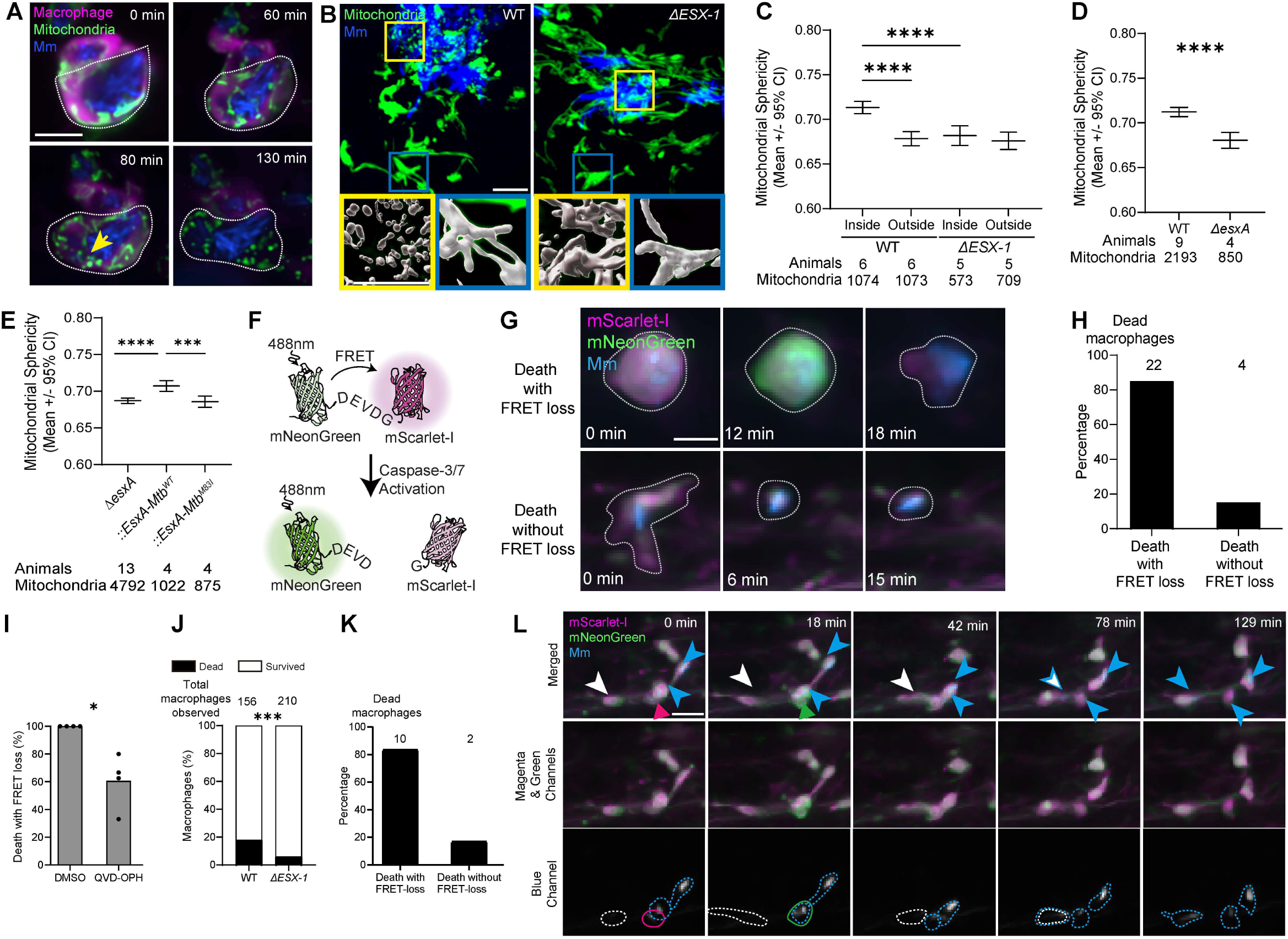
Mycobacterial EsxA induces apoptotic death of infected macrophages. (A – C and G – K) Zebrafish larvae were infected with 100 – 150 WT or 280 *ΔESX-1* Mm expressing BFP2, via the (A – C) hindbrain ventricle or (G - k caudal vein, 2 days post fertilization (dpf). (A) Time-lapse confocal micrographs of Mm-infected macrophages in an animal expressing *Tg(mfap4:MTS-EGFP:myl7:RFP)^cu72^* and *Tg(mpeg1:Brainbow)^w201^* 2 dpi. Mm (blue), macrophage mitochondria (green), and macrophages (magenta) are shown. Arrowhead, fragmented mitochondria. Scale bar, 10 μm. (B) Confocal micrograph of granulomas in *Tg(mfap4:MTS-EGFP)* animals infected with WT or *ΔESX-1* 2 dpi, showing mitochondrial morphology in macrophages inside and outside of the granuloma. Macrophage mitochondria (green) and Mm (blue). Zoomed-in boxed regions highlight differences in mitochondrial morphology between macrophages outside (blue boxes) and inside (yellow boxes) the granuloma. The lower panels show 3D renderings of the mitochondrial fluorescence signal. Scale bar, 10 μm. (C) Quantification of mitochondrial sphericity in macrophages inside and outside granulomas in *Tg(mfap4:MTS-EGFP)* animals 2 dpi. Horizontal lines indicate pooled mean values. Error bars represent 95% confidence intervals. (D and E) Zebrafish larvae were infected with ∼400 WT or ∼800 *ΔesxA* Mm expressing tdTomato via the hindbrain ventricle 2 dpf. Quantification of average mitochondrial sphericity in macrophages of *Tg(mfap4:MTS-EGFP)* animals infected with (D) WT and *ΔesxA* Mm, and (E) *ΔesxA* Mm or *ΔexsA* Mm complemented with WT or M83I point mutant Mtb *esxA*. Horizontal lines indicate pooled mean values. Error bars represent 95% confidence intervals. (F) Illustration of the FRET reporter to indicate caspases 3/7 activation. mNeonGreen (FRET donor) and mScarlet-I (FRET acceptor) fluorescent proteins are covalently linked with a peptide containing the DEVD caspase 3/7 recognition sequence. Intramolecular FRET is lost in the presence of active caspase-3 or −7. (G) Time-lapse confocal micrographs of Mm-infected macrophages in *Tg(mfap4:mNeonGreen-DEVD-mScarlet-i)* showing macrophage death with (top) and without (bottom) caspase 3/7 activation 2 days post infection (dpf). Mm (blue) and macrophages (magenta or green) are shown. Scale bar, 10 μm. (H) Quantification of macrophage death modes in wild-type animals expressing *Tg(mfap4:mNeonGreen-DEVD-mScarlet-i)* and infected with BFP2-expressing WT Mm 2 dpi. (I) Quantification of macrophage deaths at 2 dpi associated with FRET loss in animals infected intravenously with BFP2-expressing WT Mm and treated with QVD-OPH (50 μM) or 0.5% DMSO immediately following infection. (J) Quantification of observed macrophage death in wild-type animals expressing *Tg(mfap4:mNeonGreen-DEVD-mScarlet-i)* and infected with WT or *ΔESX-1* Mm 2 dpi. (K) Quantification of macrophage death modes in wild-type animals expressing *Tg(mfap4:mNeonGreen-DEVD-mScarlet-i)* and infected with BFP2-expressing *ΔESX-1* Mm 2 dpi. (L) Time-lapse confocal micrographs of a dying infected macrophage being phagocytosed by nearby macrophages in an *Tg(mfap4: mNeonGreen-DEVD-mScarlet-i)* animal 2 dpi. Mm (blue), live macrophage (magenta), dying macrophage with caspase 3/7 activation (green). Arrows and solid polygons, the dying macrophage before (magenta) and after (green) caspase 3/7 activation. Arrowheads and dashed polygons, uninfected (white), being infected (white and blue), and infected (blue) macrophages that phagocytose the dying macrophages. Scale bar, 20 μm. See Movie S1. Statistical significance was determined by (C and E) one-way ANOVA with Tukey post-test, (D and I) two-tailed, unpaired t-test, and (J) Fisher’s exact test. *\* p < 0.05*, ** *p < 0.01*, *** *p < 0.001*, and **** *p < 0.0001*. (A – D, F – K) (A and G – K), Data are representative of two or more independent experiments.

*ΔESX-1* Mm induced less mitochondrial sphericity, even in like-sized *ΔESX-1* granulomas achieved by increasing *ΔESX-1* infection inoculum size (Figure 1B, right and 1C). ESX-1’s main virulence phenotypes including membranolytic activity and cell death have been ascribed to EsxA, its major secreted substrate.^23,31–33^ We found that EsxA is responsible for ESX-1 mitochondrial damage also. *ΔesxA* infection induced less mitochondrial damage, similar to *ΔESX-1* infection (Figure 1D). However, EsxA deletion also affects secretion of other ESX-1 substrates.^34–36^ To rigorously implicate EsxA, we took advantage of an EsxA C-terminal point mutation (M83I) that abrogates membranolytic activity and macrophage death without abolishing secretion of EsxA or other ESX-1 substrates.^23,37^ Infection with *ΔesxA* Mm complemented with Mtb WT EsxA resulted in WT levels of mitochondrial damage whereas complementation with EsxA^M83I^ produced attenuated mitochondrial damage similar to *ΔESX-1* and *ΔesxA* infection (Figure 1E). These results confirmed that EsxA is responsible for ESX-1 macrophage mitochondrial damage (Figures 1B – 1E).

Next, for dynamic monitoring of ESX-1/EsxA mediated apoptosis, we engineered zebrafish to express a FRET (Förster Resonance Energy Transfer) reporter of caspase-3/7 activity under a macrophage-specific promoter. In this construct, the mNeonGreen FRET donor and mScarlet-I FRET acceptor are covalently linked with a peptide containing a caspase-3/7 recognition sequence.^38,39^ In the absence of caspase-3/7 activity, excitation of mNeonGreen with 488nm light produces green plus FRET-induced red fluorescence (shown in magenta) (Figure 1F and 1G). In the presence of active caspase-3/7 the linker is cleaved, abolishing FRET-based activation of mScarlet-I and concomitantly boosting mNeonGreen fluorescence (Figure 1F and G). We assessed death of individual macrophages over 6 hours following WT Mm infection.

85% of dying macrophages (as determined by membrane loss) had preceding caspase 3/7 activation (Figure 1G and 1H). Treatment of infected animals with the pan-caspase inhibitor QVD-OPH reduced death events associated with FRET loss by 40%, confirming that caspase activity was responsible for linker cleavage (Figure 1I). Finally, as expected, *ΔESX-1* infection had 3-fold fewer death events in the same interval and the few deaths observed were also associated with FRET loss (Figure 1J and 1K). Together, these experiments provided molecular confirmation that infected granuloma macrophages die by apoptosis, that ESX-1/EsxA promotes this death and showed the feasibility of dynamic monitoring of the death events.

Importantly, the caspase 3/7 FRET reporter could also be used to dynamically monitor the fate of the dying cells. A cardinal feature of apoptosis is that the dying cells are rapidly cleared by phagocytes through a process known as efferocytosis.^24^ Differential interference contrast imaging of zebrafish larval granulomas had shown that dying mycobacterium-infected macrophages were also engulfed by newly arriving macrophages, which became infected in the process.^3^ Using the FRET reporter, we found that of 55 infected macrophages monitored during a 6-hour time-lapse video, 8 underwent apoptosis (i.e., activated caspase 3/7) and all were efferocytosed (Figure 1L and Movie S1). The median time between caspase 3/7 activation and efferocytosis was 15 minutes (range 6 – 24 minutes). Importantly, all were engulfed before they lost plasma membrane integrity so that the mycobacteria were retained within the engulfed macrophages. In sum, the use of these two reporters has allowed us to capture, in real-time, the sequence of events mediating the ESX-1/EsxA-mediated spread of mycobacteria into new macrophages in enlarging granulomas.

### Acceleration of EsxA-mediated mitochondrial damage and apoptosis results in secondary necrosis

In adult zebrafish, as granulomas continue to enlarge, their cores become necrotic with exuberant mycobacterial growth in the resultant cellular debris.^14,19^ This granuloma breakdown also requires ESX-1/EsxA: *ΔESX-1* granulomas remain cellular even when large and, as a consequence, are paucibacillary.^14^ Thus, paradoxically, ESX-1/EsxA first builds the granuloma and then breaks it down. What is the basis of this apparent dichotomy in EsxA function?

An insight into this came from our recent finding that in mTOR-deficient zebrafish, ESX-1/EsxA promotes accelerated granuloma necrosis so that it is apparent even in the smaller granulomas of larvae.^40^ mTOR-deficient granulomas rapidly lose cellularity in an ESX-1/EsxA-dependent fashion.^40^ Lacking the mitochondrial reporter zebrafish line at the time, we had shown that cultured mTOR-deficient macrophages experience accelerated death, which is preceded by mitochondrial damage as evidenced by loss of mitochondrial membrane potential, mitochondrial fragmentation, and release of cytochrome c.^40^ As in vivo, these phenotypes are ESX-1/EsxA-dependent.^40^ Now, using the mitochondrial reporter we were able to extend these findings in vivo (Figure 2A and 2B). Using the caspase 3/7 reporter line, we confirmed that the rate of death of mTOR-deficient macrophages was higher (Figure 2C).^40^ Importantly, we found caspase-3/7 activation immediately preceded mTOR-deficient macrophage death as it did in mTOR-competent macrophage death (Figure 2D). Thus, mTOR deficiency sensitizes macrophages to EsxA-mediated mitochondrial damage and apoptosis.

**Figure 2.**
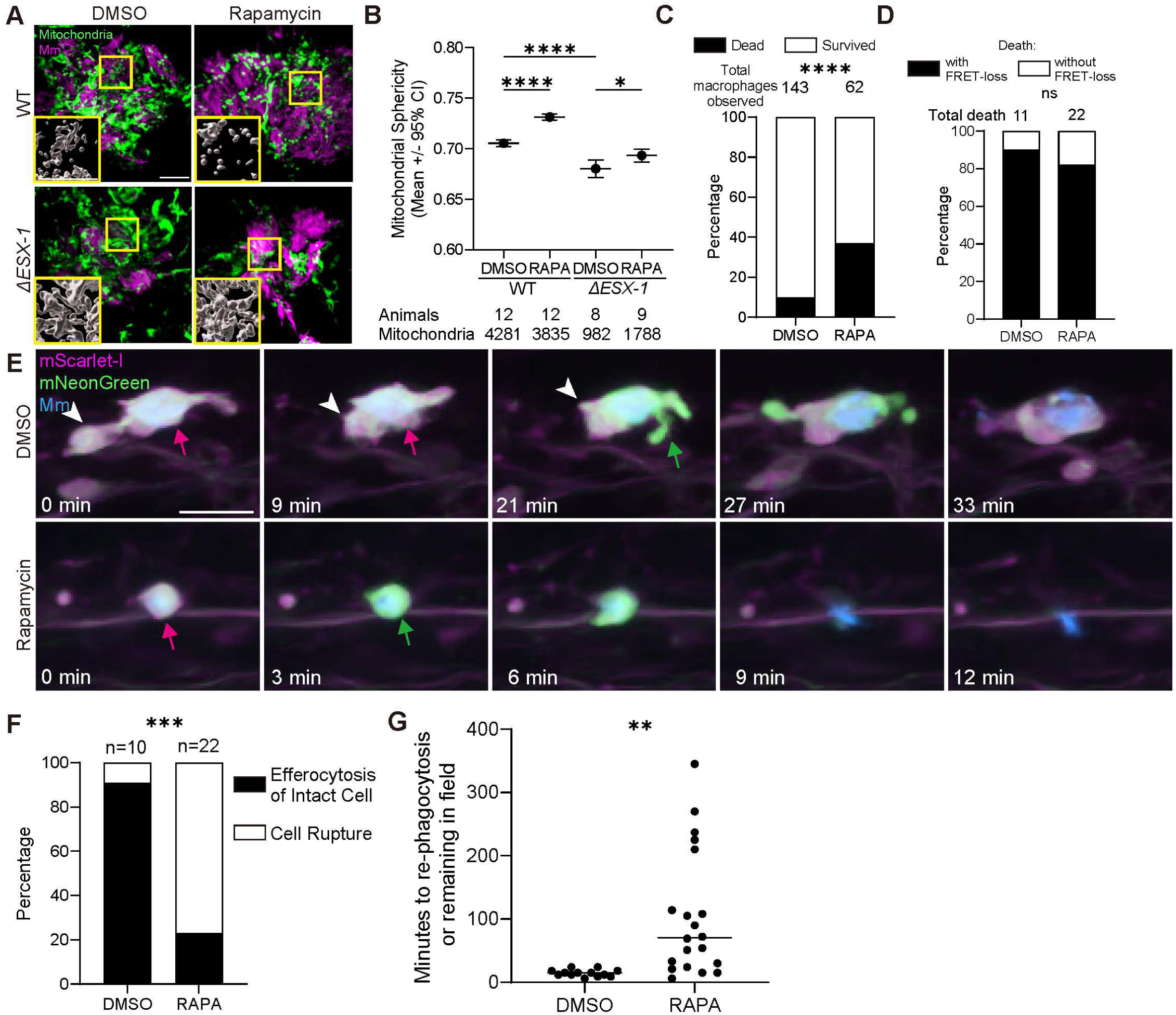
Acceleration of EsxA-induced apoptosis decimates the macrophage pool, delaying efferocytosis and causing unengulfed corpses to undergo secondary necrosis. 2 dpf zebrafish larvae were infected with 100 – 150 WT or 250 – 300 *ΔESX-1* BFP2-expressing Mm and then treated with rapamycin (1 μM) or 0.5% DMSO (vehicle control) for the duration of the experiment. (A) Confocal micrographs of granulomas in *Tg(mfap4:MTS-EGFP)* animals 2 dpi, showing mitochondrial fragmentation in the rapamycin treated, WT Mm infected group. Mm (magenta) and macrophage mitochondria (green) are shown. Zoomed-in boxed regions show 3D renderings of the mitochondrial fluorescence signal. Scale bar, 10 μm. (B) Quantification of average mitochondrial sphericity in *Tg(mfap4:MTS-EGFP)* animals 2 dpi. Horizontal lines indicate pooled mean values. Error bars represent 95% confidence intervals. (C) Quantification of macrophage deaths through 6-hour timelapse imaging in *Tg(mfap4:mNeonGreen-DEVD-mScarlet-i)* animals infected with WT Mm 3 dpi. (D) Quantification of macrophage death with or without caspases 3/7 activation through 6-hour timelapse imaging in *Tg(mfap4:mNeonGreen-DEVD-mScarlet-i)* animals infected with WT Mm 3 dpi. (E) Time-lapse confocal micrographs of apoptotic WT Mm-infected macrophages undergoing efferocytosis (top) or secondary necrosis (bottom) in a *Tg(mfap4:mNeonGreen-DEVD-mScarlet-i)* animal 3 dpi. Apoptotic macrophage before (magenta arrow) and after (green arrow) caspase-3/7 activation, efferocytic macrophage (white arrowhead). Scale bar, 20 μm. (F) Quantification of macrophage death followed by efferocytosis or secondary necrosis in *Tg(mfap4:mNeonGreen-DEVD-mScarlet-i)* animals 3 dpi. (G) Quantification of the time interval between an infected macrophage dying and being re-phagocytosed in *Tg(mfap4:mNeonGreen-DEVD-mScarlet-i)* animals 3 dpi. Symbols represent individual Mm-infected, dying macrophages. Horizontal lines indicate mean values. Statistical significance was determined by (B) one-way ANOVA with Tukey post-test, (C, D, and F) Fisher’s exact test, and (G) two-tailed, unpaired t-test. ns *p > 0.05,* ** *p < 0.01*, *** *p < 0.001*, and **** *p < 0.0001*. Data are representative of two independent experiments.

Why ESX-1/EsxA-mediated apoptosis results in necrotic breakdown of granulomas in mTOR-deficient animals became clear when we examined the fate of the dying cells. We observed that in contrast to dying WT macrophages, mTOR-deficient macrophages more frequently failed to be efferocytosed and underwent membrane loss, i.e., secondary necrosis (Figure 2E and F). This was reflected in the delayed kinetics of efferocytosis of mTOR-deficient macrophages (Figure 2G). These results lead to the conclusion that accelerated ESX-1/EsxA mediated apoptosis depletes the local supply of uninfected macrophages so that efferocytosis cannot keep pace. The unengulfed apoptotic macrophages then suffer membrane loss, releasing mycobacteria into the accumulating necrotic debris.

### HIF-1 induction in mycobacterium-infected macrophages sensitizes them to EsxA-mediated mitochondrial damage and apoptotic death

Was the early breakdown of small mTOR-deficient granulomas related to the central necrosis seen in large mTOR-competent granulomas? Two sets of clues suggested that it was. First, we had found that mTOR deficiency sensitized macrophages to ESX-1/EsxA-dependent death through impairments of mitochondrial respiration.^40^ mTOR deficiency impairs glycolytic fueling of mitochondrial energy metabolism (OXPHOS).^40^ Indeed, reducing OXPHOS through distinct means – inhibiting pyruvate entry into the Krebs cycle or genetically impairing electron transport chain (ETC) complex I assembly – accelerated ESX-1/EsxA-dependent granuloma breakdown.^40^ Second, the necrotic cores of granulomas in humans, nonhuman primates, guinea pigs, rabbits, and zebrafish experience hypoxia, which impairs mitochondrial respiration.^41–47^ Hypoxia induces the transcription factor Hypoxia Inducible Factor (HIF-1), a master regulator of host response to oxygen sensing that mediates a shift from oxidative to anaerobic respiration, and human and zebrafish tuberculous granulomas express HIF-1 or its targets.^16,44,47–51^

We hypothesized that HIF-1 induction in the macrophages in the enlarging granuloma cores might accelerate their death by sensitizing their mitochondria to ESX-1/EsxA-mediated damage. To test this, we first asked if HIF-1 activity is elevated in association with the necrotic cores of adult zebrafish granulomas. We infected transgenic adult zebrafish expressing mCherry under the transcriptional control of a HIF-response element (HRE)^52^ and micro-dissected their granulomas two weeks later. HIF-1 activity was greatest in the macrophages adjacent to and within the necrotic core (Figure 3A and Movie S2). This observation was consistent with HIF-1 accumulation in enlarging granulomas driving their necrosis.

**Figure 3.**
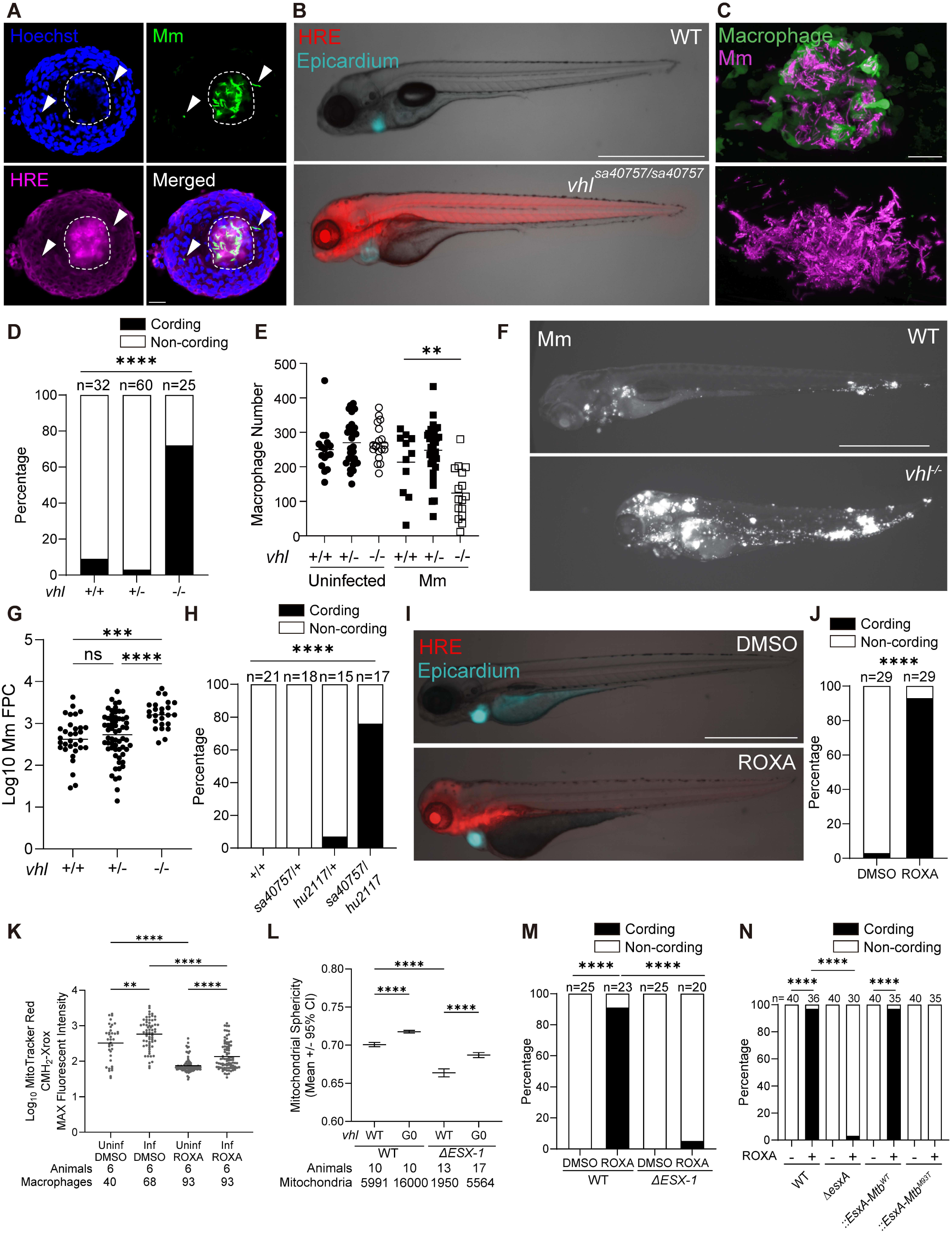
HIF-1 induction sensitizes macrophages to EsxA-mediated mitochondrial damage and increases host susceptibility to Mm infection. (A) Confocal micrograph of a granuloma with central necrosis isolated 4 weeks post infection (wpi) from a *Tg(4xhre-tata:mCherry,cmlc2:eGFP)* adult animal infected intraperitoneally with mWasabi-expressing Mm. Infected cells in non-necrotic regions of the granuloma (arrowheads) show low HIF activity compared to cells within the necrotic core (dashed lines). Hoechst-labelled nuclei (blue), Mm (green), HRE-driven mCherry expression (magenta). Scale bar, 10 μm. See Movie S2. (B) Overlaid widefield and bright field micrographs of *vhl^sa40757/sa40757^* and wild-type siblings expressing *Tg(4xhre-tata:mCherry,cmlc2:EGFP)* 5 dpf. HRE-driven mCherry expression (red), constitutive epicardial GFP signal confirms transgene expression (blue). Scale bar, 1000 μm. Zebrafish were infected via the caudal vein (C – H, J, M and N) or the hindbrain ventricle (L) with tdTomato-expressing Mm of the indicated strains 2 dpf. (C) Confocal micrographs of WT (top) and *vhl ^sa40757/sa40757^* (bottom) *Tg(mpeg1:YFP)* animals 5 dpi. Scale bar, 50 μm. (D) Quantification of extracellular mycobacterial growth (cording) in animals from vhl^sa40757/+^ incross 5dpi. (E) Quantification of macrophages in Mm-infected and age-matched uninfected animals from *vhl^sa40757/+^* incross expressing *Tg(mpeg1:YFP)* 5 dpi. Symbols represent individual animals. Horizontal lines indicate mean values. (F) Widefield micrograph of Mm fluorescence, in *vhl^sa40757/sa40757^* and WT sibling 5dpi. Scale bar, 1000 μm. (G) Quantification of Mm fluorescent pixel counts (FPC) in animals from *vhl^sa40757/+^* incross 5dpi. Symbols represent individual animals. Horizontal lines indicate mean values. (H) Quantification of mycobacterial cording 5 dpi. (I) Overlaid widefield and bright field micrographs of *Tg(4xhre-tata:mCherry,cmlc2:EGFP)* animals 5 dpf, three days after treatment with roxadustat (ROXA, 60 μM) or 0.5% DMSO. Scale bar 1000 μm. (J) Quantification of Mm cording in roxadustat (ROXA)- and vehicle-treated animals 5dpi. (K) Quantification of MitoTracker CMH_2_-Xros mean fluorescence intensity (MFI) in uninfected (bystander) and Mm-infected macrophages from *Tg(mpeg1:YFP)* animals treated with roxadustat (ROXA) or vehicle, 24 hours-post caudal vein infection with BFP2-expressing Mm and drug treatment. Symbols indicate values from individual macrophages. Horizontal lines indicate mean values. (L) Quantification of average mitochondrial sphericity in granuloma macrophages from WT and *vhl* G_0_ crispants expressing *Tg(mfap4:MTS-EGFP)* 2 dpi. Horizontal lines indicate pooled mean values. Error bars represent 95% confidence intervals. (M) Quantification of Mm cording in WT animals infected with WT or Δ*ESX-1* Mm and treated with roxadustat (ROXA) or vehicle 5dpi. (N) Mycobacterial cording in roxadustat (ROXA)- and vehicle-treated animals infected with *ΔesxA* Mm or *ΔexsA* Mm complemented with WT or point mutant Mtb *esxA*, 5dpi. Statistical significance was determined by (D, H, J, M, and N) Fisher exact test, and (E, G, K, and L) one-way ANOVA with Tukey post-test. ns *p > 0.05*, * *p < 0.05*, *** *p < 0.001*, and **** *p < 0.0001*. Data are representative of two (H and L) or three (D, G, J, K and M) independent experiments.

To test this rigorously, we used zebrafish with high HIF-1 activity created by a mutation in the von Hippel-Lindau protein (VHL). HIF-1 comprises a cytosolic oxygen-regulated unstable alpha subunit and a nuclear, constitutive beta subunit.^50^ HIF-1 activity is regulated through the binding of HIF-1α to VHL, which targets it for proteasomal degradation.^53^ VHL mutations cause HIF-1 accumulation by preventing its degradation.^53^ Therefore, VHL mutant animals should have high HIF-1 activity even in the absence of hypoxia. We confirmed that zebrafish larvae homozygous for a nonsense *vhl* mutation (*vhl^sa40757^*) had increased HIF-1 activity (HRE fluorescence) and target gene expression and exhibited the craniofacial defects, increased angiogenesis, and gasping that have been observed in other models of HIF-1 stabilization^54–57^ (Figure 3B and S1A – S1C and Movie S3). As expected, VHL mutants did not survive beyond 8 – 10 days post fertilization (dpf).

At 5 dpi, granulomas in WT fish were cellular, with mycobacteria confined to their macrophages (Figure 3C). In contrast, granulomas in VHL mutants had depleted their macrophages, resulting in greatly increased growth of the mycobacteria extracellularly in characteristic cords (Figure 3C).^4,5^ Bacterial cording is a binary phenotype, which serves as a reliable proxy for macrophage death (Figure 3D).^4,5^ We confirmed that whereas uninfected VHL mutants had similar numbers of macrophages as wildtype, infected ones had depleted their macrophages consistent with increased death (Figure 3E). As expected, the release of mycobacteria into the granulomas’ growth-permissive extracellular debris resulted in increased overall bacterial burdens in the animals (Figure 3F and 3G). We confirmed the causal role of the *vhl* mutation (as opposed to a background mutation) by crossing *vhl^sa40757^* heterozygotes to animals carrying a distinct *vhl* mutation (*hu2117*)^58^ and showing that the compound heterozygotes also had increased cording (Figure 3H). Moreover, the drug roxadustat, which increases HIF-1 activity in humans by inhibiting prolyl hydroxylases^59^, did so in zebrafish and rendered infected macrophages hyper-susceptible to mycobacterium-mediated death (Figure 3I, 3J, and S2A). Finally, the increased death of HIF-1-high macrophages was not due to their failure to control intracellular Mm burdens; WT and HIF-1-high macrophages, including dying ones, had similar intracellular burdens (Figure S2A – S2C).

To determine if mitochondrial function was impaired in HIF-1-high macrophages, we first visualized mROS production in infected and uninfected macrophages using the fluorogenic probe MitoTracker Red CM-H_2_-Xros.^60^ Elevated HIF activity impaired Mm infection-induced mROS production similarly to mTOR deficiency, which is indicative of a low OXPHOS state (Figure 3K).^40^ Next, using the transgenic macrophage mitochondrial reporter fish line, we found that the HIF-1-high state exhibited increased mitochondrial damage upon Mm infection, again similar to mTOR-deficient animals (Figure 3L). As with mTOR deficiency, ESX-1/EsxA was responsible for the increased death of HIF-1-high macrophages: HIF-1-high animals had increased bacterial cording upon wildtype but not *ΔESX-1* nor *ΔesxA* infections (Figure 3M and 3N). Moreover, complementation of the Mm *ΔesxA* strain with WT Mtb EsxA but not with EsxA with an attenuating C-terminal point mutation restored the increased cording phenotype of HIF-1-high animals (Figure 3N). Together, these experiments showed that high HIF-1 activity in granuloma macrophages sensitizes them to ESX-1/EsxA-mediated mitochondrial damage and death.

### HIF-1-high apoptotic macrophages more frequently undergo secondary necrosis

We asked if the accelerated rate of apoptosis in HIF-1-high macrophages culminated in their secondary necrosis. We first confirmed that the susceptibility of HIF-1-high macrophages to ESX-1/EsxA was cell intrinsic. We created a transgenic fish line expressing a macrophage-specific dominant active form of HIF-1, which cannot be degraded by VHL. The zebrafish genome has two *HIF1A* paralogues, *hif1aa* and *hif1ab*, but zebrafish macrophages only express *hif1ab*.^51,61^ Therefore, we created a stable transgenic line expressing dominant-active *hif1ab* under the control of the *mfap4* macrophage-specific promoter (*mfap4:DAhif-1ab*) and crossed it to a green fluorescent version of the HRE transgenic line to monitor HIF-1 activity.^52,62,63^ Macrophage-specific expression of DAhif1 induced HIF-1 activity and importantly correspondingly increased EsxA-mediated macrophage death (Figures 4A and 4B). Thus, elevated macrophage-intrinsic HIF-1 activity sensitizes the mycobacterium-infected macrophage to EsxA-mediated macrophage death.

**Figure 4.**
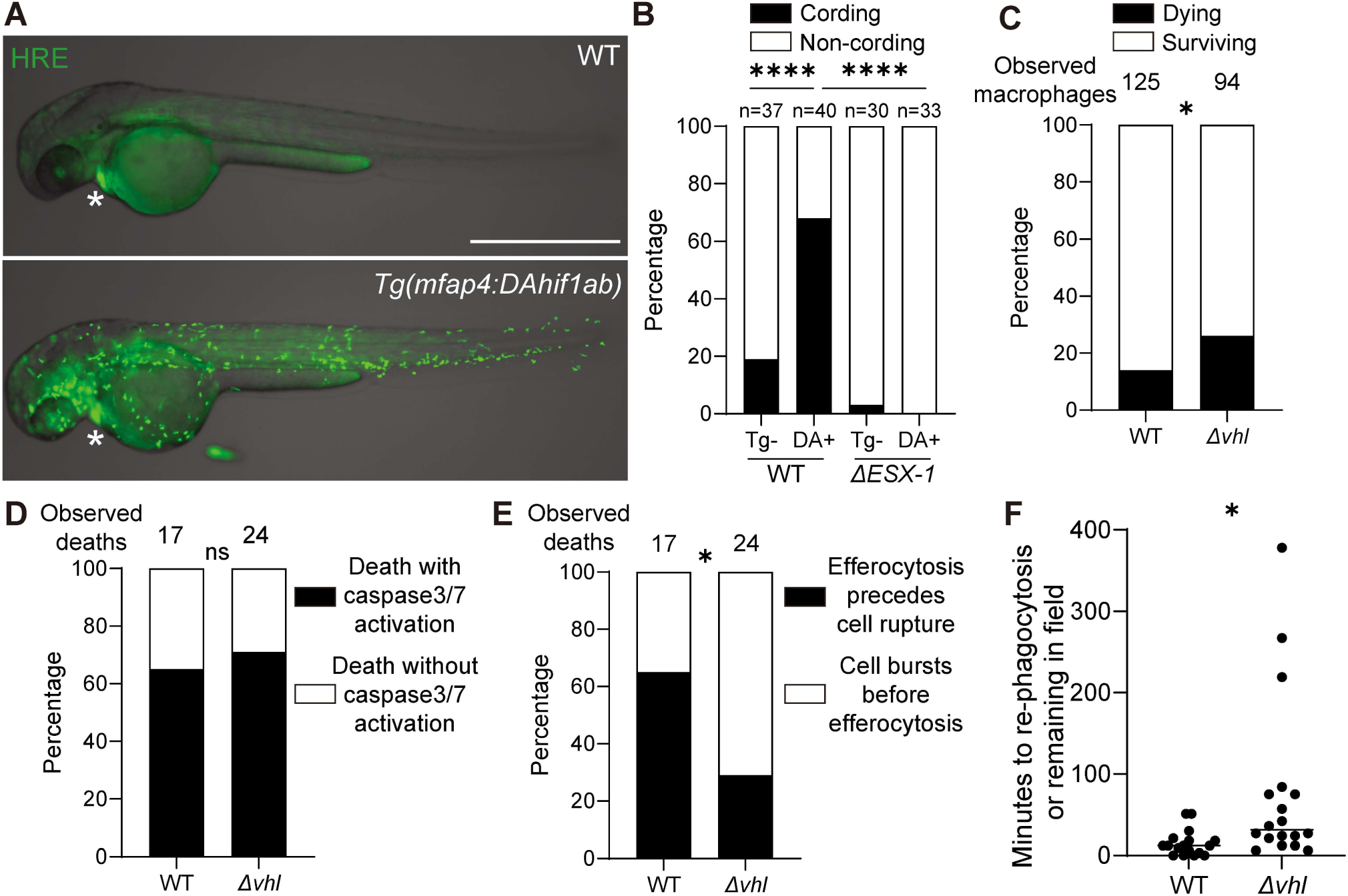
HIF-1 accelerates ESX-1/EsxA-mediated apoptosis to instigate the secondary necrosis of unengulfed macrophage corpses. (A) Overlaid widefield and bright field micrographs of *Tg(mfap4:DAhif1ab)* animals, which specifically express dominant-active HIF-1 in macrophages, and WT siblings expressing *Tg(4xhre-tata:EGFP,cmlc2:EGFP)* 3 dpf. Epicardial GFP expression (asterisk). Scale bar 1000 μm. (B) Quantification of mycobacterial cording 5dpi. (C – F) WT and *vhl* G_0_ crispants expressing *Tg(mfap4:mNeonGreen-DEVD-mScarlet-i)* were infected with ∼100 BFP2-expressing WT Mm and imaged by timelapse microscopy for 6 hrs. See Movie S3. (C) Quantification of macrophage deaths 3 dpi. (D) Quantification of macrophage deaths with or without caspases 3/7 activation in wild-type or *vhl* G_0_ crispants expressing *Tg(mfap4:mNeonGreen-DEVD-mScarlet-i)* with BFP2-expressing WT Mm 3 dpi.(E) Quantification of macrophage deaths followed by their efferocytosis or secondary necrosis in wild-type or *vhl* G_0_ crispants expressing *Tg(mfap4:mNeonGreen-DEVD-mScarlet-i)* 3 dpi. (F) Quantification of the time interval from infected macrophages undergoing cell death to Mm getting re-phagocytosed in wild-type or *vhl* G_0_ crispants expressing *Tg(mfap4:mNeonGreen-DEVD-mScarlet-i)* 3 dpi. Symbols represent infected macrophage death. Horizontal bars indicate median values. Statistical significance was determined by (B - E) Fisher exact test or (F) two-tailed, unpaired t-test. ns *p > 0.05*, * *p < 0.05*, and **** *p < 0.0001*. Data are representative of two (C-F) or more (B) independent experiments.

Next, using the caspase 3/7 reporter line, we confirmed that more infected HIF-high macrophages died and saw that most death events were preceded by caspase 3/7 activation (Figure 4C and 4D). Finally, we showed that the kinetics of efferocytosis were delayed leading to a higher frequency of secondary necrosis (Figure 4E and 4F). Thus, as with mTOR deficiency, the increased rate of EsxA-mediated apoptosis of HIF-1-high macrophages outpaces their efferocytosis, leading to secondary necrosis and mycobacterial release into the extracellular milieu.

## DISCUSSION

This work reveals how a single mycobacterial virulence factor, EsxA, orchestrates cellular granuloma enlargement and then its necrosis through a single mechanism – namely mitochondrial damage of infected macrophages leading to apoptotic death. EsxA is reported to also trigger nonapoptotic modes of cell death, such as pyroptosis and necrosis induced by lipid mediators or type I interferons, including through neutrophil-mediated immunopathology.^28,64–68^ This work shows that these death modes need not be invoked to explain why the cores of large granulomas invariably break down.^9^ The switch from granuloma enlargement to its necrotic breakdown can be explained by the accelerated kinetics of EsxA-mediated mitochondrial damage in response to HIF-1 activation in the hypoxic centers of enlarging granulomas. Thus, mycobacteria induce a host metabolic adaptation to hypoxia and exploit this adaptation to choreograph the steps needed for transmission - granuloma enlargement followed by its breakdown.

HIF-1 increases macrophage mycobactericidal activity in the context of interferon-γ (IFNγ)-mediated CD4^+^ T cell immunity though not in the sole context of innate immunity.^69^ Therefore, as adaptive immunity comes into play, the macrophages’ enhanced antimicrobial capacity might be expected to offset their increased vulnerability to EsxA-mediated mitochondrial damage and cytotoxicity. However, infected macrophages deep in the granuloma interior have limited access to CD4^+^ T cell-derived IFNγ. This impairment is because Mtb-specific CD4^+^ T cells must physically interact with infected macrophages to enhance their mycobactericidal activity and the T cells primarily localize to the granuloma rim.^70–72^ The few CD4^+^ T cells that do migrate into granulomas tend to produce little IFNγ compared to those outside them.^71,73,74^ These factors may convert the granuloma core, where HIF-1 is stabilized, into an area where only innate immunity is operant. Consistent with this idea, Mtb burdens are elevated in the T cell-poor, hypoxic zones of nonhuman primate TB granulomas relative to the non-hypoxic zones.^75^ Thus, even in the context of the adaptive immune phase of infection, the lack of access of T cells to the granuloma core can make it vulnerable to EsxA–HIF-1 necrosis.

This work finds that the ultimate cause of the necrotic phenotype is the depletion of the local supply of efferocytic macrophages that would keep mycobacteria intracellular. On the one hand, efferocytosis promotes mycobacterial expansion through spread into new macrophages in the enlarging granuloma.^3^ On the other hand, it can restrict burdens by retaining mycobacteria intracellularly, as evidenced both by conditions where the supply of efferocytic macrophages is diminished^76,77^ or, as shown in this study, where macrophage apoptosis is accelerated. This is true in the context of fully mature epithelioid granulomas seen in adult zebrafish.^76^ Indeed, the fact that efferocytosis prevents mycobacterial release into the growth-promoting extracellular space may explain the paradoxical observations that efferocytosis reduces mycobacterial loads while simultaneously increasing the number of macrophages that become infected.^3,78,79^

Because of the importance of granuloma necrosis to disease progression and transmission, modes of programmed macrophage necrosis in TB are widely researched.^8,80–82^ These include diverse mechanisms that accelerate necrosis of mycobacterium-infected macrophages. ^5,40,60,68,77,83–91^ This work reveals the default mode by which enlarging granulomas progress to necrosis, even in the absence of these programmed necrosis modes – through the interaction of an apoptosis-inducing mycobacterial virulence factor with an ordinarily host-protective determinant.

## RESOURCE AVAILABILITY

### Lead contact

Requests for resources, reagents, and additional information should be directed to and will be fulfilled by the lead contact, Lalita Ramakrishnan.

### Materials availability

Plasmids and zebrafish lines generated in this study can be obtained from the lead contact.

### Data and code availability

-Microscopy data reported in this work will be shared by the lead contact upon request.

-This paper does not report any original code.

-Any additional information needed to re-analyze the data reported in this work will be provided by the lead contact upon request.

## Supporting information

Resources Table

## ACKNOWLEDGEMENTS

This work was supported by grants from the National Institutes of Health (MERIT award R37-AI054503, 5R01-AI054503) and a Wellcome Trust Principal Research Fellowship (223103/Z/21/Z) (L.R.); a Gates Cambridge Scholarship (OPP1144)(B.L.); a Human Frontier Science Program Postdoctoral Fellowship (LT0029/2022-L), a Blavatnik Cambridge Postdoctoral Fellowship, a Rothschild Yad Hanadiv Fellowship and the Israel National Postdoctoral Award for Advancing Women in Science (M.E.B.); and an Esther Ehrmann Lazard Faculty Scholar appointment (A.J.P.).

We thank L. Lee for contributing experiments and ideas; J. Milão and R. Santhakumar for technical assistance; F. Roca and K. Shanahan for insights and discussion; P. Edelstein for guidance and help on statistical analysis; M. Behr, B. Cormack, P. Edelstein, and D. Tobin for manuscript review; P. Elks and A. Vettori for sharing plasmids; N. Goodwin and the University of Cambridge aquatics facility staff for zebrafish husbandry; the LMB’s light microscopy core for access to microscopes and analysis software; and the LMB’s media service for preparation of reagents for bacterial and zebrafish culture.

## AUTHOR CONTRIBUTIONS

A.J.P. and L.R. conceived and oversaw the project. All authors designed experiments and analyzed data. A.J.P., B.L., and M.E.B. did experiments. E.M.B-N provided the *vhl* alleles from a mutagenesis screen. A.J.P., B.L., and M.E.B. made figures. A.J.P., B.L., M.E.B., and L.R. wrote the paper.

## METHODS

### Zebrafish husbandry and infections

Zebrafish of the AB wild-type (Zebrafish International Resource Center, ZIRC), TL strain (ZIRC), or of mixed AB/TL backgrounds were used in experiments. Zebrafish embryos and larvae of undetermined sex (due to their early stage of development) were used in experiments. Adult male zebrafish were used in granuloma explant experiments. Zebrafish husbandry and experiments were conducted in compliance with guidelines from the UK Home Office, using protocols approved by the Animal Welfare and Ethical Review Body of the University of Cambridge.

The *Tg(mpeg1:Brainbow)^w201^* ^76^ *Tg(mfap4:tdTomato-CAAX)^xt6^*^62^, *Tg(mfap4:MTS-EGFP;myl7:RFP)^cu72^* (this study), *Tg(mfap4:mNeonGreen-DEVD-mScarlet-I)^cu40^* (this study), *Tg(4xtata-hre:mCherry;mlc2:eGFP)^cu73^*(this study), *Tg(mfap4:DAhif1ab-2A-tdTomato-CAAX)^cu74^* (this study), Tg(*4xhre-tata:EGFP)^cu75^* (this study) and *Tg(−7kdrl:DsRed2*)*^pd27^* fluorescent reporter lines were maintained in the AB strain background. *vhl^sa40757^* ^92^and *vhl^hu2117^* ^58^ mutant lines were generated in the TL strain and maintained in mixed AB/TL backgrounds.

Zebrafish were maintained in buffered reverse osmotic water systems under a 14-hr light/10-hr dark cycle. Larvae were fed paramecia twice daily and juvenile and adult zebrafish were fed at least twice a day with dry food and brine shrimp. Zebrafish embryos were collected and cultured in reverse osmosis (RO) water containing 0.18 g/L Instant Ocean Salt supplemented with 0.25 mg/mL methylene blue at 28.5°C. 1 dpf embryos used in experiments were transferred to 0.5x E2 medium (7.5 mM NaCl, 0.25 mM KCl, 0.5 mM MgSO_4_, 0.075 mM KH_2_PO_4_, 0.025 mM Na_2_HPO_4_, 0.5 mM CaCl_2_, and 0.35 mM NaHCO_3_) supplemented with 0.003% PTU (1-phenyl-2-thiourea, Sigma) to prevent pigmentation.

For infections, 2 dpf larvae were dechorionated manually or with ≤ 0.5mg/ml pronase (Sigma-Aldrich) and subsequently anesthetized in fish waster containing 0.025% tricaine (Fisher Scientific). Larvae were microinjected via the hindbrain ventricle or the caudal vein using single-cell suspensions of Mm of known concentrations to deliver 100 – 250 bacteria per 3 – 5 nL injection as previously described ^93^. Phenol red sodium salt (Sigma-Aldrich) diluted in PBS (≤ 1% w/v) was used as an injection tracer. Adult zebrafish were injected intraperitoneally with a single-cell suspension of Mm to deliver 20 – 100 bacteria using a similar method to the one developed by Cosma, et al.^94^

In experiments with mutant zebrafish lines, wild-type siblings were used as controls. Animals were genotyped after the completion of the experiment. In experiments involving drug treatments, infected larvae were randomly distributed between treatment groups. Drug treatment was initiated immediately after completion of the infection session. Drugs were administered through soaking. Drug treatment was maintained for the duration of the experiment.

## METHOD DETAILS

### Bacterial strains

*M. marinum* M strain (ATCC #BAA-535) and its mutant derivatives Δ*ESX-1* and Δ*esxA* were grown in Middlebrook 7H9 medium (BD Difco) supplemented with glycerol, oleic acid, albumin, dextrose, and Tween-80 (Sigma-Aldrich)^23^ Mm strains transformed with plasmids expressing BFP2, mWasabi, or tdTomato under the control of the *msp12* promoter or complementation constructs encoding wild-type Mtb or EsxA point mutations were grown under hygromycin B (Cambridge Bioscience) or kanamycin (Sigma-Aldrich) selection, as appropriate.

### Creation of transgenic lines

The plasmid used to generate the *Tg(mfap4:MTS-EGFP:myl7:RFP)^cu72^* was created by cloning EGFP tagged with a mitochondrial targeting sequence (MTS-EGFP) from (RRID:Addgene_31241)^95^ into a pCS2-based Tol2 vector containing the *mfap4* macrophage-specific promoter sequence and the myl7 promoter sequence driving RFP expression in the heart^84^, using the Gibson Cloning kit (New England Biolabs). The plasmid used to generate the *Tg(mfap4:mNeonGreen-DEVD-mScarlet-I)^cu40^* was assembled from a PCR fragments containing the FRET donor mNeonGreen coding sequence plus an 18-amino acid peptide containing the caspase-3/7 recognition sequence (underlined) SSSELSG<u>DEVD</u>GTSGSEF^38^, the mScarlet-I coding sequence^39^, and the pCS2-based Tol2 vector containing the *mfap4* macrophage-specific promoter sequence^62^ using the Gibson Cloning kit (New England Biolabs). *Tg(4xtata-hre:mCherry;mlc2:eGFP)^cu73^* and Tg(*4xhre-tata:EGFP)*^cu75^ were created by injecting plasmids provided by Dr. Andrea Vettori.^52^ The plasmid to create *Tg(mfap4:DAhif1ab-2A-tdTomato-CAAX)^cu74^* (DAhif) line assembled from fragments of *pUAS:da-hif-1*α*b-IRES-GFP*^96^, provided by Dr. Phil Elks and pTol2-mfap4:nlsVenus-2A-tdTomato-CAAX using the Gibson Cloning kit (New England Biolabs). Dominant-active HIF1ab protein contains three point mutations (P402A, P564G, N804A) which prevent its hydroxylation by prolyl hydroxylases (PHDs) and Factor-inhibiting hypoxia-inducible factor (FIH).^63^

Correct assembly of each plasmid was confirmed by diagnostic PCRs of the joined segments and by Sanger sequencing. Tol2 transposase RNA (RRID:Addgene_51818)^97^ was in vitro transcribed with the T7 mMessage/mMachine kit (Thermo Fisher) according to the manufacturer’s instructions. Plasmids and transposase RNA were diluted in Tango Buffer (Thermo Scientific) containing 2% phenol red sodium salt solution (Sigma) and injected into one-cell stage embryos.^98^ G0 transgenic larvae were identified by florescence microscopy and were raised to adulthood. A single F1 founder fish identified through pairwise crosses of G0 adults and non-transgenic AB wild-type fish was used to establish each transgenic line.

### CRISPR-Cas9 mutagenesis

The mutagenesis procedure was adapted from published methods^99^ and performed as previously described.^40^ *vhl* G0 CRISPR mutants (crispants) were created with a pool of three guide RNAs targeting exon 1 of *vhl*. Crispants in the B cell-specific gene *aicda*, which encodes Activation-induced Cytidine Deaminase, were used as negative controls. AID is almost exclusively expressed by B cells undergoing immunoglobulin class-switch recombination in mammals and affinity maturation in both mammals and fish and is not expressed by zebrafish phagocytes or larvae.^100^

To create guide RNAs (60 µM), Alt-R tracrRNA and individual target-specific Alt-R crRNAs (*vhl*: Dr.Cas9.VHL.1.AA, AB, AC; IDT*; aicda*: Dr.Cas9.AICDA.1.AA; IDT) were mixed at an equimolar ratio in nuclease-free Duplex Buffer (IDT) and incubated at 95°C for 5 minutes. Aliquots of complexed RNA were stored at −20°C. Ribonucleoprotein complexes (RNPs) were produced by mixing total complexed RNA and Alt-R Sp Cas9 Nuclease V3 (IDT) at an equimolar ratio (e.g., 0.31 pmol of total RNA and 0.31 of pmol Cas9 protein) in working buffer (20 mM HEPES, 150 mM KCl, pH 7.5) and incubated at 37°C for 10 minutes. 1-cell stage zebrafish embryos were injected with ∼2 nL of RNPs. Successful mutagenesis was determined by High Resolution Melt Analysis (HRMA)^102^of PCR products produced with primers spanning the crRNA target sites.

### Zebrafish genotyping

DNA from adult fin clips or whole larvae was extracted using the HotSHOT method.^101^ Fish were genotyped by HRMA^102^ or Kompetitive Allele-Specific PCR (KASP, LGC Biosearch) of PCR products (see Key Resources Table), using a CFX Connect or CFX Duet Real-time PCR instruments (BioRad).

### Mm infections

For larval infections, single-cell suspensions of Mm were mixed with phenol red sodium salt (1% w/v diluted in PBS, Sigma-Aldrich) and used to deliver a known titer of bacteria in a 3 – 5 nL injection via the caudal vein or the hindbrain ventricle, as previously described. ^93^ Infections were performed on 2 dpf larvae which were dechorionated manually or with 0.5% mg/mL pronase (Sigma-Aldrich) and then anesthetized in fish water containing 0.025% tricaine (Sigma).

For adult infections, fish were anesthetized in aquatics system water containing 0.02% tricaine and immediately injected intraperitoneally with a single-cell suspension of Mm. A 10 µL microliter syringe affixed with a 26G needle (Hamilton) was used to deliver 20 – 100 Mm in 5 μL of PBS.

For each experiment, injection inoculums were determined by injecting the same volume onto bacteriological plates (Middlebrook 7H10 agar plates supplemented with glycerol, oleic acid, albumin, dextrose, and hygromycin B or kanamycin, as appropriate). Mm colonies were counted after 10 – 14 days of growth in a 33°C incubator.

### Granuloma explants

Granulomas from adult zebrafish were collected 4 weeks post infection as previously described.^15^ Zebrafish were euthanized with a tricaine overdose, surface decontaminated with 70% ethanol, and placed on ice-cold PBS. Carcasses were transferred to glass petri dishes containing L-15 media (Thermo Fisher) and dissected under a stereomicroscope. Explanted granulomas were washed three times in L-15 media and fixed overnight in 4% paraformaldehyde (PFA) solution (4% w/v PFA in PBS, Alfa Aesar). To label nuclei, explanted granulomas were stained with Hoechst 33342 (Tocris) diluted in PBS. Granulomas were transferred to optical bottom plates (MatTek) for fluorescence microscopy.

### mROS detection

To measure mROS production in zebrafish macrophages, larvae were injected with 3 – 5 nL of 10 mM MitoTracker Red CMXH_2_-Xros in PBS via the caudal vein immediately prior to imaging by confocal microscopy.^85^

### Microscopy and image analyses

In most experiments, larvae were anesthetized in PTU 0.5x E2 medium containing 0.025% tricaine. In experiments involving serial imaging of infections, larvae were cryo-anesthetized on ice for 10 min. as previously described.^93^ Transgenic larvae were manually sorted under widefield fluorescence illumination on a Nikon SMZ18 stereomicroscope or an upright Nikon E600 compound light microscope. To assess relative bacterial burdens and mycobacterial cording, larvae were imaged on a Nikon Eclipse Ti-E inverted microscope using 4x or 10x objectives. Relative bacterial burdens were determined by quantifying fluorescent pixel counts with an Image J macro (National Institutes of Health) on manually processed samples ^11^ or with an automated image analysis processing tool. ^103^

For confocal microscopy, larvae were embedded in 1% low melt agarose (TopVision) on glass-bottom plates (MatTek or WillCo-dish). Laser scanning confocal microscopy was performed on a Nikon A1R confocal microscope with 20x (0.75 NA Air) or 40x (1 NA Water) objectives and the galvano or resonant scanners as previously described.^40^ 0.3 - 2 μm optical sections were combined to generate 15 - 80 μm z stacks. Similar settings were used to image explanted granulomas. Time-lapse experiments were performed at 27°C using a microscope incubator (Okolab) with acquisition intervals of 2 - 5 minutes for 6 hours. Images were processed using the denoising feature in the Elements software (Nikon). Spinning disk confocal microscopy was performed using a Nikon W1 Spinning Disk inverted microscope with a 20x (0.75NA Air) or 60x (1.2NA Water) objectives and a sCMOS camera (95% QE). 0.3 - 2 μm optical sections were combined to generate 15 - 100 μm z stacks. Time-lapse microscopy was performed at room temperature using acquisition intervals of 2 – 10 minutes for 4 – 8 hours.

Confocal micrographs were processed with ImageJ or Imaris (Bitplane Scientific Software). Mm volume and mitochondrial sphericity were determined using the Imaris surface rendering feature. Bacteria inside granulomas were manually identified. Then, mitochondria within 5 µm from granuloma bacteria were defined as mitochondria within the granuloma. This distance threshold was defined by inspecting representative granuloma micrographs where a cytosolic macrophage fluorescent marker was used to define granuloma boundaries.

Macrophage cell death in timelapse experiments with *Tg(mfap4:mNeonGreen-DEVD-mScarlet-I)^cu40^* animals was defined by the sudden loss of mNeonGreen fluorescence. Apoptotic macrophages were identified by the sequential loss of FRET-dependent mScarlet-I fluorescence (indicative of caspase-3/7 activation) and mNeonGreen fluorescence. Macrophages undergoing non-apoptotic death modes were identified by the simultaneous loss of mNeonGreen and mScarlet-I fluorescence. To monitor the efferocytosis and secondary necrosis of infected apoptotic macrophages, co-localized fluorescence signals from the Mm and macrophage reporters were tracked manually frame-by-frame. Re-phagocytosis kinetics were assessed by recording the time interval between macrophage death and re-phagocytosis of corpse-associated mycobacteria by nearby macrophages or the time corpse-associated Mm remained in the field of view until the end of the movie.

### Reverse Transcription Quantitative Polymerase Chain Reaction (RT-qPCR) assays

Total RNA from biological replicates consisting of 6 pooled larvae was isolated with Qiagen RNeasy kit. RNA samples also underwent an on-column DNAse I digest procedure (Qiagen) to remove any contaminating DNA. cDNA was produced with a cDNA synthesis kit (Takara) using Superscript II reverse transcriptase and oligo DT primers. qPCRs were performed on biological duplicates or triplicates using the 2x SYBR Green PCR Master Mix (Applied Biosystems) and run on a CFX Connect thermocycler (Bio-Rad). Values from technical duplicates were averaged and used to calculate fold changes in target gene expression with the 2^-ΔΔCT^ method, using *ef1a* as a reference gene.

### Quantification and statistical analyses

The following statistical analyses were performed in Prism (GraphPad): unpaired, two-tailed Student’s t test (used to determine statistical significance between the mean values of two groups); One-way Analysis of Variance (ANOVA) with Tukey’s post-test (used to compare the mean values of three or more groups); and Fisher’s exact test (used to analyze contingency table data). Horizontal lines represent means or medians. p values are as follows: ns, not significant, * p < 0.05; ** p < 0.01; *** p < 0.001; **** p < 0.0001. The statistical tests used for each figure can be found in the corresponding figure legend. n represents the number of animals per experimental group.

---

**Movie S1. Death of Mm-infected macrophages, related to Figure 1**.

Time-lapse confocal microscopy of a dying infected macrophage being phagocytosed by nearby live macrophages in an *Tg(mfap4: mNeonGreen-DEVD-mScarlet-i)* animal 2dpi. Mm (blue), live macrophage (magenta), dying macrophage with caspase 3/7 activation (green). Arrowheads, the dying macrophage before (yellow) and after (green) caspase 3/7 activation phagocytosed by uninfected (white) and infected (blue) macrophages. Scale bar, 20 μm.

**Movie S2. HIF transcriptional activity is induced in Mm granulomas of adult zebrafish, related to Figure 3**.

Optical sections of confocal micrographs showing an explant granuloma with central necrosis isolated 4 weeks post infection (wpi) from a *Tg(4xhre-tata:mCherry,cmlc2:eGFP)* adult animal infected intraperitoneally with mWasabi-expressing Mm. Hoechst-labelled nuclei (blue), Mm (green), HRE-driven mCherry expression (magenta). Scale bar, 10 μm.

**Movie S3. *vhl* mutant zebrafish larvae display defects in craniofacial developmental and increased gasping, related to Figure 3**.

Timelapse brightfield microscopy of a *vhl* homozygous mutant zebrafish larva and a phenotypically WT sibling.

**Figure S1.**
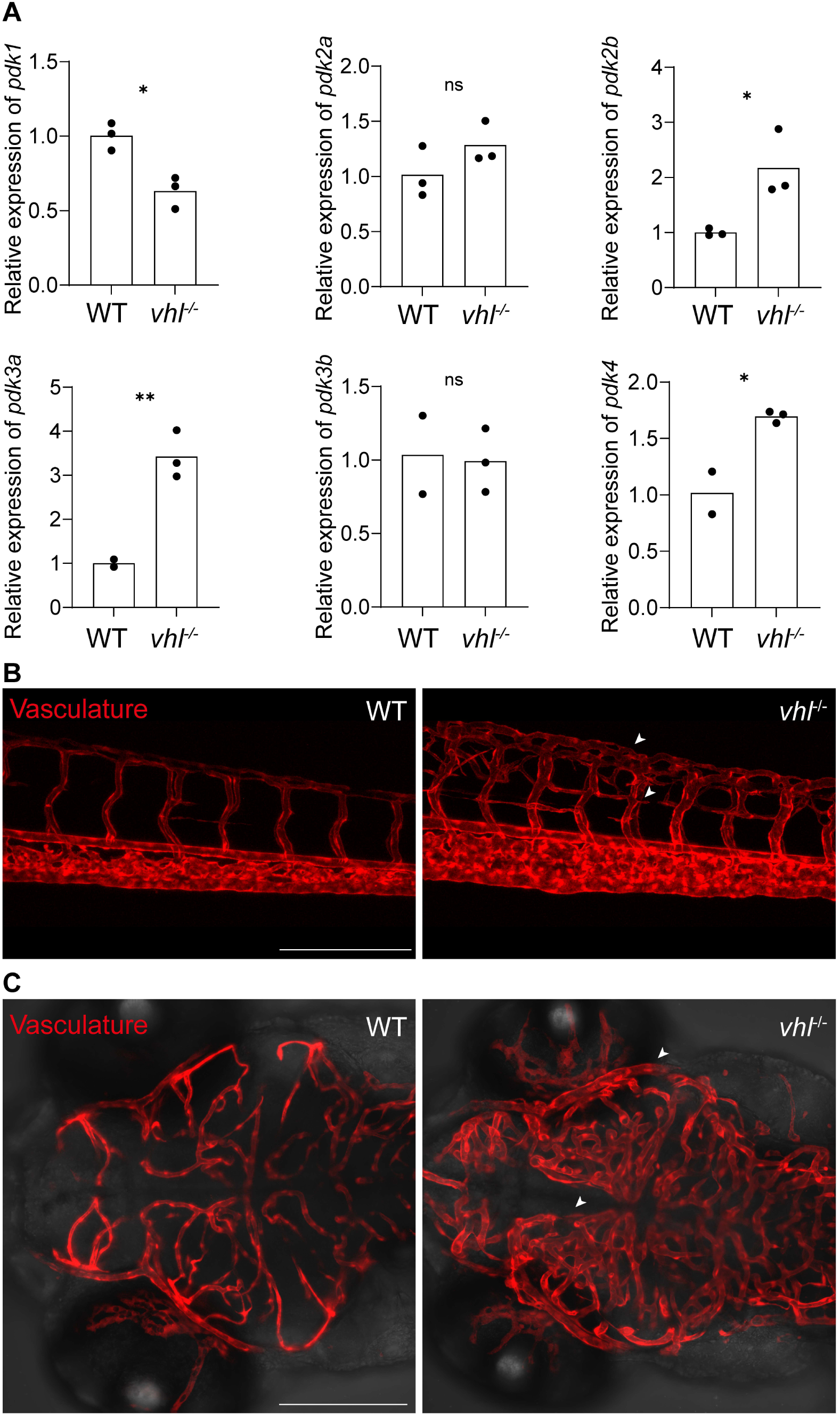
*vhl* mutant animals have increased HIF-1 activity, related to Figure 3. (A) Expression of HIF target gene transcripts in uninfected *vhl^sa40757/sa40757^* and phenotypically WT siblings (*vhl^+/+^* and *vhl^+/-^*) 3dpf, measured by qRT-PCR and normalized to wild-type values. (B and C) Confocal micrographs of the (B) tail or (C) brain vasculature of *vhl^sa40757/sa40757^* and phenotypically WT sibling, 5dpf. Arrowheads indicate increased blood vessel abundance. Scale bar, 100 μm.

**Figure S2.**
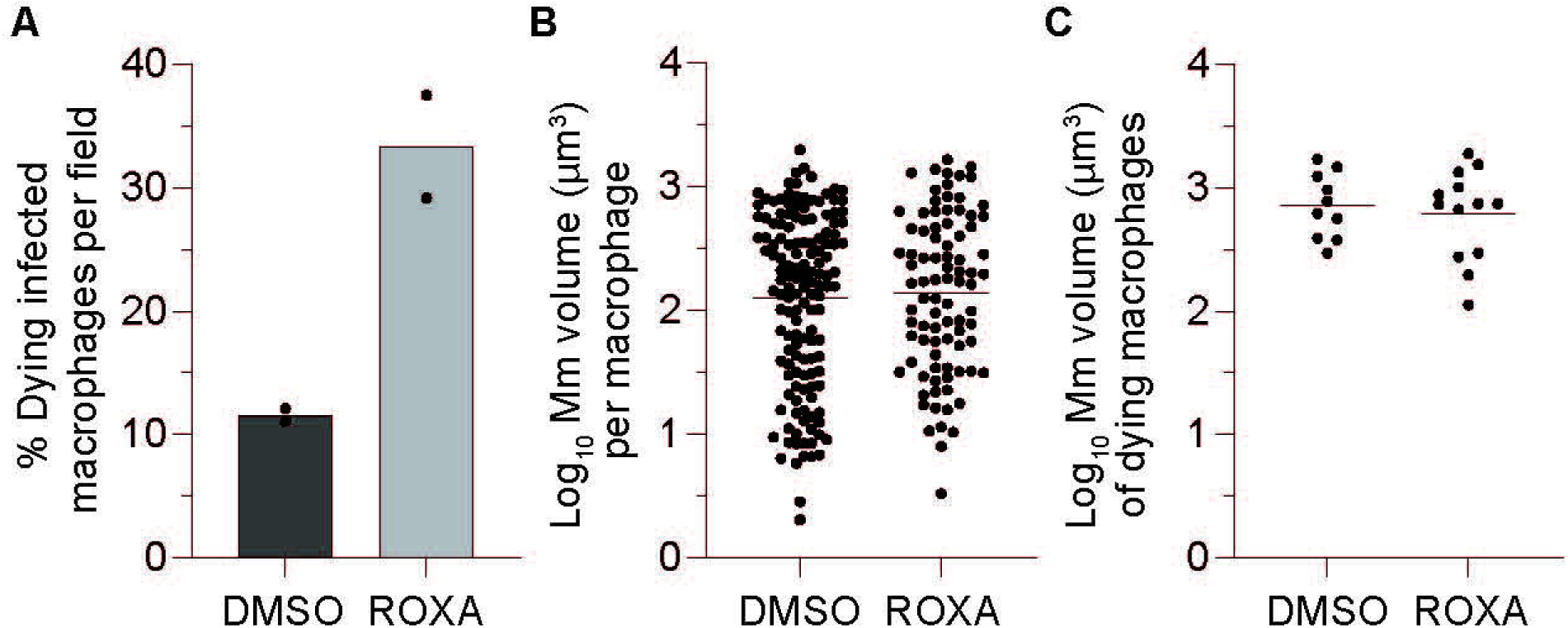
HIF-1 stabilization accelerates Mm-induced macrophage cell death without altering intracellular Mm burdens, related to Figure 3. *Tg(mpeg1;YFP)* larvae were infected intravenously with tdTomato-expressing Mm at 2 dpf and treated with Roxadustat (60 μM) or 0.5% DMSO immediately after infection. 5 dpf (3 dpi) animals were imaged by timelapse microscopy for 4 hours. Dying macrophages were identified by rapid loss of YFP fluorescence accompanied by cellular fragmentation. (A) Percent of infected macrophages dying per field. (B) Intracellular Mm burdens per macrophage in the first movie frame. (C) Intracellular Mm burdens of dying macrophages at the time of cell death. Bars and horizontal lines indicate mean values. Symbols represent individual (A) animals or (B and (C) macrophages. (B and C) Statistical significance was determined with a two-tailed, unpaired t*-*test with Welch’s correction (ns, *p > 0.05*).

## REFERENCES

1. Pagan, A.J., and Ramakrishnan, L. (2018). The Formation and Function of Granulomas. Annu Rev Immunol 36, 639–665. 10.1146/annurev-immunol-032712-100022.

2. Ramakrishnan, L. (2012). Revisiting the role of the granuloma in tuberculosis. Nat Rev Immunol 12, 352–366. 10.1038/nri3211.

3. Davis, J.M., and Ramakrishnan, L. (2009). The role of the granuloma in expansion and dissemination of early tuberculous infection. Cell 136, 37–49. 10.1016/j.cell.2008.11.014.

4. Clay, H., Volkman, H.E., and Ramakrishnan, L. (2008). Tumor necrosis factor signaling mediates resistance to mycobacteria by inhibiting bacterial growth and macrophage death. Immunity 29, 283–294. 10.1016/j.immuni.2008.06.011.

5. Tobin, D.M., Vary, J.C., Jr., Ray, J.P., Walsh, G.S., Dunstan, S.J., Bang, N.D., Hagge, D.A., Khadge, S., King, M.C., Hawn, T.R., et al. (2010). The lta4h locus modulates susceptibility to mycobacterial infection in zebrafish and humans. Cell 140, 717–730. 10.1016/j.cell.2010.02.013.

6. Kramnik, I., Dietrich, W.F., Demant, P., and Bloom, B.R. (2000). Genetic control of resistance to experimental infection with virulent Mycobacterium tuberculosis. Proc Natl Acad Sci U S A 97, 8560–8565. 10.1073/pnas.150227197.

7. Russell, D.G., Cardona, P.J., Kim, M.J., Allain, S., and Altare, F. (2009). Foamy macrophages and the progression of the human tuberculosis granuloma. Nat Immunol 10, 943–948. 10.1038/ni.1781.

8. Pagan, A.J., and Ramakrishnan, L. (2014). Immunity and Immunopathology in the Tuberculous Granuloma. Cold Spring Harb Perspect Med 5. 10.1101/cshperspect.a018499.

9. Urbanowski, M.E., Ordonez, A.A., Ruiz-Bedoya, C.A., Jain, S.K., and Bishai, W.R. (2020). Cavitary tuberculosis: the gateway of disease transmission. Lancet Infect Dis 20, e117–e128. 10.1016/S1473-3099(20)30148-1.

10. Ong, C.W., Elkington, P.T., and Friedland, J.S. (2014). Tuberculosis, pulmonary cavitation, and matrix metalloproteinases. Am J Respir Crit Care Med 190, 9–18. 10.1164/rccm.201311-2106PP.

11. Ramakrishnan, L. (2013). The zebrafish guide to tuberculosis immunity and treatment. Cold Spring Harb Symp Quant Biol 78, 179–192. 10.1101/sqb.2013.78.023283.

12. Canetti, G. (1955). The tubercle bacillus in the pulmonary lesion of man; histobacteriology and its bearing on the therapy of pulmonary tuberculosis, American rev. ed. Edition (New York, Springer).

13. Rich, A. (1951). The Pathogenesis of Tuberculosis. 2nd ed. (Charles C Thomas).

14. Swaim, L.E., Connolly, L.E., Volkman, H.E., Humbert, O., Born, D.E., and Ramakrishnan, L. (2006). Mycobacterium marinum infection of adult zebrafish causes caseating granulomatous tuberculosis and is moderated by adaptive immunity. Infect Immun 74, 6108–6117. 10.1128/IAI.00887-06.

15. Cronan, M.R., Matty, M.A., Rosenberg, A.F., Blanc, L., Pyle, C.J., Espenschied, S.T., Rawls, J.F., Dartois, V., and Tobin, D.M. (2018). An explant technique for high-resolution imaging and manipulation of mycobacterial granulomas. Nat Methods 15, 1098–1107. 10.1038/s41592-018-0215-8.

16. Viswanathan, G., Hughes, E.J., Gan, M., Xet-Mull, A.M., Lowy, J.P., Pyle, C.J., Alexander, G., Swain-Lenz, D., Liu, Q., and Tobin, D.M. (2026). Granuloma dual RNA-seq reveals composite transcriptional programs driven by neutrophils and necrosis within tuberculous granulomas. Sci Adv 12, eadw4619. 10.1126/sciadv.adw4619.

17. Pozos, T.C., and Ramakrishnan, L. (2004). New models for the study of Mycobacterium-host interactions. Curr Opin Immunol 16, 499–505. 10.1016/j.coi.2004.05.011.

18. Davis, J.M., Clay, H., Lewis, J.L., Ghori, N., Herbomel, P., and Ramakrishnan, L. (2002). Real-time visualization of mycobacterium-macrophage interactions leading to initiation of granuloma formation in zebrafish embryos. Immunity 17, 693–702. 10.1016/s1074-7613(02)00475-2.

19. Volkman, H.E., Clay, H., Beery, D., Chang, J.C., Sherman, D.R., and Ramakrishnan, L. (2004). Tuberculous granuloma formation is enhanced by a mycobacterium virulence determinant. PLoS Biol 2, e367. 10.1371/journal.pbio.0020367.

20. Volkman, H.E., Pozos, T.C., Zheng, J., Davis, J.M., Rawls, J.F., and Ramakrishnan, L. (2010). Tuberculous granuloma induction via interaction of a bacterial secreted protein with host epithelium. Science 327, 466–469. 10.1126/science.1179663.

21. Cambier, C.J., Takaki, K.K., Larson, R.P., Hernandez, R.E., Tobin, D.M., Urdahl, K.B., Cosma, C.L., and Ramakrishnan, L. (2014). Mycobacteria manipulate macrophage recruitment through coordinated use of membrane lipids. Nature 505, 218–222. 10.1038/nature12799.

22. Cambier, C.J., O’Leary, S.M., O’Sullivan, M.P., Keane, J., and Ramakrishnan, L. (2017). Phenolic Glycolipid Facilitates Mycobacterial Escape from Microbicidal Tissue-Resident Macrophages. Immunity 47, 552–565 e554. 10.1016/j.immuni.2017.08.003.

23. Osman, M.M., Shanahan, J.K., Chu, F., Takaki, K.K., Pinckert, M.L., Pagan, A.J., Brosch, R., Conrad, W.H., and Ramakrishnan, L. (2022). The C terminus of the mycobacterium ESX-1 secretion system substrate ESAT-6 is required for phagosomal membrane damage and virulence. Proc Natl Acad Sci U S A 119, e2122161119. 10.1073/pnas.2122161119.

24. Morioka, S., Maueroder, C., and Ravichandran, K.S. (2019). Living on the Edge: Efferocytosis at the Interface of Homeostasis and Pathology. Immunity 50, 1149–1162. 10.1016/j.immuni.2019.04.018.

25. Whittaker, J.A., Bentley, D.P., Melville-Jones, G.R., and Slater, A.J. (1976). Granuloma formation in patients receiving BCG immunotherapy. J Clin Pathol 29, 693–697. 10.1136/jcp.29.8.693.

26. Shea, C.R., Imber, M.J., Cropley, T.G., Cosimi, A.B., and Sober, A.J. (1989). Granulomatous eruption after BCG vaccine immunotherapy for malignant melanoma. J Am Acad Dermatol 21, 1119–1122. 10.1016/s0190-9622(89)70310-8.

27. Narita, K., Akita, H., Kikuchi, E., Nakahara, T., Okuda, S., Nakatsuka, S., Oya, M., and Jinzaki, M. (2019). Biopsy-diagnosed renal granuloma after intravesical bacillus Calmette-Guerin therapy for bladder carcinoma: a case series and review of the literature. BJR Case Rep 5. 10.1259/bjrcr.20190012.

28. Gern, B.H., Klas, J.M., Foster, K.A., Kanagy, M.E., Cohen, S.B., Plumlee, C.R., Duffy, F.J., Neal, M.L., Halima, M., Gustin, A.T., et al. (2025). Early and opposing neutrophil and CD4 T cell responses shape pulmonary tuberculosis pathology. J Exp Med 222. 10.1084/jem.20250161.

29. Lienard, J., Nobs, E., Lovins, V., Movert, E., Valfridsson, C., and Carlsson, F. (2020). The Mycobacterium marinum ESX-1 system mediates phagosomal permeabilization and type I interferon production via separable mechanisms. Proc Natl Acad Sci U S A 117, 1160–1166. 10.1073/pnas.1911646117.

30. Wiens, K.E., and Ernst, J.D. (2016). The Mechanism for Type I Interferon Induction by Mycobacterium tuberculosis is Bacterial Strain-Dependent. PLoS Pathog 12, e1005809. 10.1371/journal.ppat.1005809.

31. Zhang, Q., Wang, D., Jiang, G., Liu, W., Deng, Q., Li, X., Qian, W., Ouellet, H., and Sun, J. (2016). EsxA membrane-permeabilizing activity plays a key role in mycobacterial cytosolic translocation and virulence: effects of single-residue mutations at glutamine 5. Sci Rep 6, 32618. 10.1038/srep32618.

32. Augenstreich, J., Haanappel, E., Sayes, F., Simeone, R., Guillet, V., Mazeres, S., Chalut, C., Mourey, L., Brosch, R., Guilhot, C., and Astarie-Dequeker, C. (2020). Phthiocerol Dimycocerosates From Mycobacterium tuberculosis Increase the Membrane Activity of Bacterial Effectors and Host Receptors. Front Cell Infect Microbiol 10, 420. 10.3389/fcimb.2020.00420.

33. Bates, T.A., Trank-Greene, M., Wrynla, X.H., Anastas, A., Gurmessa, S.K., Merutka, I.R., Dixon, S.D., Shumate, A., Groncki, A.R., Parson, M.A.H., et al. (2024). ESAT-6 undergoes self-association at phagosomal pH and an ESAT-6-specific nanobody restricts M. tuberculosis growth in macrophages. Elife 12. 10.7554/eLife.91930.

34. Fortune, S.M., Jaeger, A., Sarracino, D.A., Chase, M.R., Sassetti, C.M., Sherman, D.R., Bloom, B.R., and Rubin, E.J. (2005). Mutually dependent secretion of proteins required for mycobacterial virulence. Proc Natl Acad Sci U S A 102, 10676–10681. 10.1073/pnas.0504922102.

35. Champion, M.M., Williams, E.A., Pinapati, R.S., and Champion, P.A. (2014). Correlation of phenotypic profiles using targeted proteomics identifies mycobacterial esx-1 substrates. J Proteome Res 13, 5151–5164. 10.1021/pr500484w.

36. Bao, Y., Wang, L., and Sun, J. (2021). A Small Protein but with Diverse Roles: A Review of EsxA in Mycobacterium-Host Interaction. Cells 10. 10.3390/cells10071645.

37. Brodin, P., de Jonge, M.I., Majlessi, L., Leclerc, C., Nilges, M., Cole, S.T., and Brosch, R. (2005). Functional analysis of early secreted antigenic target-6, the dominant T-cell antigen of Mycobacterium tuberculosis, reveals key residues involved in secretion, complex formation, virulence, and immunogenicity. J Biol Chem 280, 33953–33959. 10.1074/jbc.M503515200.

38. Mayer, C.T., Gazumyan, A., Kara, E.E., Gitlin, A.D., Golijanin, J., Viant, C., Pai, J., Oliveira, T.Y., Wang, Q., Escolano, A., et al. (2017). The microanatomic segregation of selection by apoptosis in the germinal center. Science 358. 10.1126/science.aao2602.

39. McCullock, T.W., MacLean, D.M., and Kammermeier, P.J. (2020). Comparing the performance of mScarlet-I, mRuby3, and mCherry as FRET acceptors for mNeonGreen. PLoS One 15, e0219886. 10.1371/journal.pone.0219886.

40. Pagan, A.J., Lee, L.J., Edwards-Hicks, J., Moens, C.B., Tobin, D.M., Busch-Nentwich, E.M., Pearce, E.L., and Ramakrishnan, L. (2022). mTOR-regulated mitochondrial metabolism limits mycobacterium-induced cytotoxicity. Cell 185, 3720–3738 e3713. 10.1016/j.cell.2022.08.018.

41. Via, L.E., Lin, P.L., Ray, S.M., Carrillo, J., Allen, S.S., Eum, S.Y., Taylor, K., Klein, E., Manjunatha, U., Gonzales, J., et al. (2008). Tuberculous granulomas are hypoxic in guinea pigs, rabbits, and nonhuman primates. Infect Immun 76, 2333–2340. 10.1128/IAI.01515-07.

42. Harper, J., Skerry, C., Davis, S.L., Tasneen, R., Weir, M., Kramnik, I., Bishai, W.R., Pomper, M.G., Nuermberger, E.L., and Jain, S.K. (2012). Mouse model of necrotic tuberculosis granulomas develops hypoxic lesions. J Infect Dis 205, 595–602. 10.1093/infdis/jir786.

43. Oehlers, S.H., Cronan, M.R., Scott, N.R., Thomas, M.I., Okuda, K.S., Walton, E.M., Beerman, R.W., Crosier, P.S., and Tobin, D.M. (2015). Interception of host angiogenic signalling limits mycobacterial growth. Nature 517, 612–615. 10.1038/nature13967.

44. Belton, M., Brilha, S., Manavaki, R., Mauri, F., Nijran, K., Hong, Y.T., Patel, N.H., Dembek, M., Tezera, L., Green, J., et al. (2016). Hypoxia and tissue destruction in pulmonary TB. Thorax 71, 1145–1153. 10.1136/thoraxjnl-2015-207402.

45. Papandreou, I., Cairns, R.A., Fontana, L., Lim, A.L., and Denko, N.C. (2006). HIF-1 mediates adaptation to hypoxia by actively downregulating mitochondrial oxygen consumption. Cell Metab 3, 187–197. 10.1016/j.cmet.2006.01.012.

46. Kim, J.W., Tchernyshyov, I., Semenza, G.L., and Dang, C.V. (2006). HIF-1-mediated expression of pyruvate dehydrogenase kinase: a metabolic switch required for cellular adaptation to hypoxia. Cell Metab 3, 177–185. 10.1016/j.cmet.2006.02.002.

47. Denko, N.C. (2008). Hypoxia, HIF1 and glucose metabolism in the solid tumour. Nat Rev Cancer 8, 705–713. 10.1038/nrc2468.

48. Palazon, A., Goldrath, A.W., Nizet, V., and Johnson, R.S. (2014). HIF transcription factors, inflammation, and immunity. Immunity 41, 518–528. 10.1016/j.immuni.2014.09.008.

49. Walmsley, S.R., Cadwallader, K.A., and Chilvers, E.R. (2005). The role of HIF-1alpha in myeloid cell inflammation. Trends Immunol 26, 434–439. 10.1016/j.it.2005.06.007.

50. Walmsley, S.R., McGovern, N.N., Whyte, M.K., and Chilvers, E.R. (2008). The HIF/VHL pathway: from oxygen sensing to innate immunity. Am J Respir Cell Mol Biol 38, 251–255. 10.1165/rcmb.2007-0331TR.

51. Cronan, M.R., Hughes, E.J., Brewer, W.J., Viswanathan, G., Hunt, E.G., Singh, B., Mehra, S., Oehlers, S.H., Gregory, S.G., Kaushal, D., and Tobin, D.M. (2021). A non-canonical type 2 immune response coordinates tuberculous granuloma formation and epithelialization. Cell 184, 1757–1774 e1714. 10.1016/j.cell.2021.02.046.

52. Vettori, A., Greenald, D., Wilson, G.K., Peron, M., Facchinello, N., Markham, E., Sinnakaruppan, M., Matthews, L.C., McKeating, J.A., Argenton, F., and van Eeden, F.J.M. (2017). Glucocorticoids promote Von Hippel Lindau degradation and Hif-1alpha stabilization. Proc Natl Acad Sci U S A 114, 9948–9953. 10.1073/pnas.1705338114.

53. Kaelin, W.G., Jr. (2022). Von Hippel-Lindau disease: insights into oxygen sensing, protein degradation, and cancer. J Clin Invest 132. 10.1172/JCI162480.

54. Hickey, M.M., Lam, J.C., Bezman, N.A., Rathmell, W.K., and Simon, M.C. (2007). von Hippel-Lindau mutation in mice recapitulates Chuvash polycythemia via hypoxia-inducible factor-2alpha signaling and splenic erythropoiesis. J Clin Invest 117, 3879–3889. 10.1172/JCI32614.

55. Hickey, M.M., Richardson, T., Wang, T., Mosqueira, M., Arguiri, E., Yu, H., Yu, Q.C., Solomides, C.C., Morrisey, E.E., Khurana, T.S., et al. (2010). The von Hippel-Lindau Chuvash mutation promotes pulmonary hypertension and fibrosis in mice. J Clin Invest 120, 827–839. 10.1172/JCI36362.

56. Metelo, A.M., Noonan, H.R., Li, X., Jin, Y., Baker, R., Kamentsky, L., Zhang, Y., van Rooijen, E., Shin, J., Carpenter, A.E., et al. (2015). Pharmacological HIF2alpha inhibition improves VHL disease-associated phenotypes in zebrafish model. J Clin Invest 125, 1987–1997. 10.1172/JCI73665.

57. Kim, H.R., Greenald, D., Vettori, A., Markham, E., Santhakumar, K., Argenton, F., and van Eeden, F. (2017). Zebrafish as a model for von Hippel Lindau and hypoxia-inducible factor signaling. Methods Cell Biol 138, 497–523. 10.1016/bs.mcb.2016.07.001.

58. van Rooijen, E., Voest, E.E., Logister, I., Korving, J., Schwerte, T., Schulte-Merker, S., Giles, R.H., and van Eeden, F.J. (2009). Zebrafish mutants in the von Hippel-Lindau tumor suppressor display a hypoxic response and recapitulate key aspects of Chuvash polycythemia. Blood 113, 6449–6460. 10.1182/blood-2008-07-167890.

59. Zhu, X., Jiang, L., Wei, X., Long, M., and Du, Y. (2022). Roxadustat: Not just for anemia. Front Pharmacol 13, 971795. 10.3389/fphar.2022.971795.

60. Roca, F.J., and Ramakrishnan, L. (2013). TNF dually mediates resistance and susceptibility to mycobacteria via mitochondrial reactive oxygen species. Cell 153, 521–534. 10.1016/j.cell.2013.03.022.

61. Athanasiadis, E.I., Botthof, J.G., Andres, H., Ferreira, L., Lio, P., and Cvejic, A. (2017). Single-cell RNA-sequencing uncovers transcriptional states and fate decisions in haematopoiesis. Nat Commun 8, 2045. 10.1038/s41467-017-02305-6.

62. Walton, E.M., Cronan, M.R., Beerman, R.W., and Tobin, D.M. (2015). The Macrophage-Specific Promoter mfap4 Allows Live, Long-Term Analysis of Macrophage Behavior during Mycobacterial Infection in Zebrafish. PLoS One 10, e0138949. 10.1371/journal.pone.0138949.

63. Elks, P.M., van Eeden, F.J., Dixon, G., Wang, X., Reyes-Aldasoro, C.C., Ingham, P.W., Whyte, M.K., Walmsley, S.R., and Renshaw, S.A. (2011). Activation of hypoxia-inducible factor-1alpha (Hif-1alpha) delays inflammation resolution by reducing neutrophil apoptosis and reverse migration in a zebrafish inflammation model. Blood 118, 712–722. 10.1182/blood-2010-12-324186.

64. Beckwith, K.S., Beckwith, M.S., Ullmann, S., Saetra, R.S., Kim, H., Marstad, A., Asberg, S.E., Strand, T.A., Haug, M., Niederweis, M., et al. (2020). Plasma membrane damage causes NLRP3 activation and pyroptosis during Mycobacterium tuberculosis infection. Nat Commun 11, 2270. 10.1038/s41467-020-16143-6.

65. Chen, M., Gan, H., and Remold, H.G. (2006). A mechanism of virulence: virulent Mycobacterium tuberculosis strain H37Rv, but not attenuated H37Ra, causes significant mitochondrial inner membrane disruption in macrophages leading to necrosis. J Immunol 176, 3707–3716. 10.4049/jimmunol.176.6.3707.

66. Chen, M., Divangahi, M., Gan, H., Shin, D.S., Hong, S., Lee, D.M., Serhan, C.N., Behar, S.M., and Remold, H.G. (2008). Lipid mediators in innate immunity against tuberculosis: opposing roles of PGE2 and LXA4 in the induction of macrophage death. J Exp Med 205, 2791–2801. 10.1084/jem.20080767.

67. Divangahi, M., Chen, M., Gan, H., Desjardins, D., Hickman, T.T., Lee, D.M., Fortune, S., Behar, S.M., and Remold, H.G. (2009). Mycobacterium tuberculosis evades macrophage defenses by inhibiting plasma membrane repair. Nat Immunol 10, 899–906. 10.1038/ni.1758.

68. Zhang, L., Jiang, X., Pfau, D., Ling, Y., and Nathan, C.F. (2021). Type I interferon signaling mediates Mycobacterium tuberculosis-induced macrophage death. J Exp Med 218. 10.1084/jem.20200887.

69. Braverman, J., Sogi, K.M., Benjamin, D., Nomura, D.K., and Stanley, S.A. (2016). HIF-1alpha Is an Essential Mediator of IFN-gamma-Dependent Immunity to Mycobacterium tuberculosis. J Immunol 197, 1287–1297. 10.4049/jimmunol.1600266.

70. Srivastava, S., and Ernst, J.D. (2013). Cutting edge: Direct recognition of infected cells by CD4 T cells is required for control of intracellular Mycobacterium tuberculosis in vivo. J Immunol 191, 1016–1020. 10.4049/jimmunol.1301236.

71. Cohen, S.B., Gern, B.H., and Urdahl, K.B. (2022). The Tuberculous Granuloma and Preexisting Immunity. Annu Rev Immunol 40, 589–614. 10.1146/annurev-immunol-093019-125148.

72. Mortensen, R., Arlehamn, C.S.L., Coler, R.N., Gerner, M.Y., Goletti, D., Lewinsohn, D.A., Modlin, R.L., Musvosvi, M., Rengarajan, J., Urdahl, K.B., et al. (2026). T cell-macrophage interactions in tuberculosis: What we’ve got here is failure to communicate. J Intern Med 299, 44–65. 10.1111/joim.70028.

73. Sallin, M.A., Kauffman, K.D., Riou, C., Du Bruyn, E., Foreman, T.W., Sakai, S., Hoft, S.G., Myers, T.G., Gardina, P.J., Sher, A., et al. (2018). Host resistance to pulmonary Mycobacterium tuberculosis infection requires CD153 expression. Nat Microbiol 3, 1198–1205. 10.1038/s41564-018-0231-6.

74. Gern, B.H., Adams, K.N., Plumlee, C.R., Stoltzfus, C.R., Shehata, L., Moguche, A.O., Busman-Sahay, K., Hansen, S.G., Axthelm, M.K., Picker, L.J., et al. (2021). TGFbeta restricts expansion, survival, and function of T cells within the tuberculous granuloma. Cell Host Microbe 29, 594–606 e596. 10.1016/j.chom.2021.02.005.

75. McCaffrey, E.F., Delmastro, A.C., Fitzhugh, I., Ranek, J.S., Douglas, S., Peters, J.M., Fullaway, C.C., Bosse, M., Liu, C.C., Gillen, C., et al. (2025). The immunometabolic topography of tuberculosis granulomas governs cellular organization and bacterial control. bioRxiv. 10.1101/2025.02.18.638923.

76. Pagan, A.J., Yang, C.T., Cameron, J., Swaim, L.E., Ellett, F., Lieschke, G.J., and Ramakrishnan, L. (2015). Myeloid Growth Factors Promote Resistance to Mycobacterial Infection by Curtailing Granuloma Necrosis through Macrophage Replenishment. Cell Host Microbe 18, 15–26. 10.1016/j.chom.2015.06.008.

77. Berg, R.D., Levitte, S., O’Sullivan, M.P., O’Leary, S.M., Cambier, C.J., Cameron, J., Takaki, K.K., Moens, C.B., Tobin, D.M., Keane, J., and Ramakrishnan, L. (2016). Lysosomal Disorders Drive Susceptibility to Tuberculosis by Compromising Macrophage Migration. Cell 165, 139–152. 10.1016/j.cell.2016.02.034.

78. Martin, C.J., Booty, M.G., Rosebrock, T.R., Nunes-Alves, C., Desjardins, D.M., Keren, I., Fortune, S.M., Remold, H.G., and Behar, S.M. (2012). Efferocytosis is an innate antibacterial mechanism. Cell Host Microbe 12, 289–300. 10.1016/j.chom.2012.06.010.

79. Zheng, W., Chang, I.C., Limberis, J., Budzik, J.M., Zha, B.S., Howard, Z., Chen, L., and Ernst, J.D. (2024). Mycobacterium tuberculosis resides in lysosome-poor monocyte-derived lung cells during chronic infection. PLoS Pathog 20, e1012205. 10.1371/journal.ppat.1012205.

80. Srinivasan, L., Ahlbrand, S., and Briken, V. (2014). Interaction of Mycobacterium tuberculosis with host cell death pathways. Cold Spring Harb Perspect Med 4. 10.1101/cshperspect.a022459.

81. Vu, A., Glassman, I., Campbell, G., Yeganyan, S., Nguyen, J., Shin, A., and Venketaraman, V. (2024). Host Cell Death and Modulation of Immune Response against Mycobacterium tuberculosis Infection. Int J Mol Sci 25. 10.3390/ijms25116255.

82. Faysal, M.A., Hanafy, M., Zinniel, D.K., Tanni, F.Y., Muthukrishnan, E., Rathnaiah, G., and Barletta, R.G. (2025). Cell death pathways in response to Mycobacterium tuberculosis and other mycobacterial infections. Infect Immun 93, e0040125. 10.1128/iai.00401-25.

83. Tobin, D.M., Roca, F.J., Oh, S.F., McFarland, R., Vickery, T.W., Ray, J.P., Ko, D.C., Zou, Y., Bang, N.D., Chau, T.T., et al. (2012). Host genotype-specific therapies can optimize the inflammatory response to mycobacterial infections. Cell 148, 434–446. 10.1016/j.cell.2011.12.023.

84. Roca, F.J., Whitworth, L.J., Redmond, S., Jones, A.A., and Ramakrishnan, L. (2019). TNF Induces Pathogenic Programmed Macrophage Necrosis in Tuberculosis through a Mitochondrial-Lysosomal-Endoplasmic Reticulum Circuit. Cell 178, 1344–1361 e1311. 10.1016/j.cell.2019.08.004.

85. Roca, F.J., Whitworth, L.J., Prag, H.A., Murphy, M.P., and Ramakrishnan, L. (2022). Tumor necrosis factor induces pathogenic mitochondrial ROS in tuberculosis through reverse electron transport. Science 376, eabh2841. 10.1126/science.abh2841.

86. Amaral, E.P., Costa, D.L., Namasivayam, S., Riteau, N., Kamenyeva, O., Mittereder, L., Mayer-Barber, K.D., Andrade, B.B., and Sher, A. (2019). A major role for ferroptosis in Mycobacterium tuberculosis-induced cell death and tissue necrosis. J Exp Med 216, 556–570. 10.1084/jem.20181776.

87. Amaral, E.P., Foreman, T.W., Namasivayam, S., Hilligan, K.L., Kauffman, K.D., Barbosa Bomfim, C.C., Costa, D.L., Barreto-Duarte, B., Gurgel-Rocha, C., Santana, M.F., et al. (2022). GPX4 regulates cellular necrosis and host resistance in Mycobacterium tuberculosis infection. J Exp Med 219. 10.1084/jem.20220504.

88. Amaral, E.P., Namasivayam, S., Queiroz, A.T.L., Fukutani, E., Hilligan, K.L., Aberman, K., Fisher, L., Bomfim, C.C.B., Kauffman, K., Buchanan, J., et al. (2024). BACH1 promotes tissue necrosis and Mycobacterium tuberculosis susceptibility. Nat Microbiol 9, 120–135. 10.1038/s41564-023-01523-7.

89. Pajuelo, D., Gonzalez-Juarbe, N., Tak, U., Sun, J., Orihuela, C.J., and Niederweis, M. (2018). NAD(+) Depletion Triggers Macrophage Necroptosis, a Cell Death Pathway Exploited by Mycobacterium tuberculosis. Cell Rep 24, 429–440. 10.1016/j.celrep.2018.06.042.

90. Lee, J., Remold, H.G., Ieong, M.H., and Kornfeld, H. (2006). Macrophage apoptosis in response to high intracellular burden of Mycobacterium tuberculosis is mediated by a novel caspase-independent pathway. J Immunol 176, 4267–4274. 10.4049/jimmunol.176.7.4267.

91. Pan, H., Yan, B.S., Rojas, M., Shebzukhov, Y.V., Zhou, H., Kobzik, L., Higgins, D.E., Daly, M.J., Bloom, B.R., and Kramnik, I. (2005). Ipr1 gene mediates innate immunity to tuberculosis. Nature 434, 767–772. 10.1038/nature03419.

92. Kettleborough, R.N., Busch-Nentwich, E.M., Harvey, S.A., Dooley, C.M., de Bruijn, E., van Eeden, F., Sealy, I., White, R.J., Herd, C., Nijman, I.J., et al. (2013). A systematic genome-wide analysis of zebrafish protein-coding gene function. Nature 496, 494–497. 10.1038/nature11992.

93. Takaki, K., Davis, J.M., Winglee, K., and Ramakrishnan, L. (2013). Evaluation of the pathogenesis and treatment of Mycobacterium marinum infection in zebrafish. Nat Protoc 8, 1114–1124. 10.1038/nprot.2013.068.

94. Cosma, C.L., Swaim, L.E., Volkman, H., Ramakrishnan, L., and Davis, J.M. (2006). Zebrafish and frog models of Mycobacterium marinum infection. Curr Protoc Microbiol Chapter 10, Unit 10B 12. 10.1002/0471729256.mc10b02s3.

95. Kim, M.J., Kang, K.H., Kim, C.H., and Choi, S.Y. (2008). Real-time imaging of mitochondria in transgenic zebrafish expressing mitochondrially targeted GFP. Biotechniques 45, 331–334. 10.2144/000112909.

96. Elks, P.M., Brizee, S., van der Vaart, M., Walmsley, S.R., van Eeden, F.J., Renshaw, S.A., and Meijer, A.H. (2013). Hypoxia inducible factor signaling modulates susceptibility to mycobacterial infection via a nitric oxide dependent mechanism. PLoS Pathog 9, e1003789. 10.1371/journal.ppat.1003789.

97. Khattak, S., Murawala, P., Andreas, H., Kappert, V., Schuez, M., Sandoval-Guzman, T., Crawford, K., and Tanaka, E.M. (2014). Optimized axolotl (Ambystoma mexicanum) husbandry, breeding, metamorphosis, transgenesis and tamoxifen-mediated recombination. Nat Protoc 9, 529–540. 10.1038/nprot.2014.040.

98. Suster, M.L., Abe, G., Schouw, A., and Kawakami, K. (2011). Transposon-mediated BAC transgenesis in zebrafish. Nat Protoc 6, 1998–2021. 10.1038/nprot.2011.416.

99. Wu, R.S., Lam, II, Clay, H., Duong, D.N., Deo, R.C., and Coughlin, S.R. (2018). A Rapid Method for Directed Gene Knockout for Screening in G0 Zebrafish. Dev Cell 46, 112–125 e114. 10.1016/j.devcel.2018.06.003.

100. White, R.J., Collins, J.E., Sealy, I.M., Wali, N., Dooley, C.M., Digby, Z., Stemple, D.L., Murphy, D.N., Billis, K., Hourlier, T., et al. (2017). A high-resolution mRNA expression time course of embryonic development in zebrafish. Elife 6. 10.7554/eLife.30860.

101. Truett, G.E., Heeger, P., Mynatt, R.L., Truett, A.A., Walker, J.A., and Warman, M.L. (2000). Preparation of PCR-quality mouse genomic DNA with hot sodium hydroxide and tris (HotSHOT). Biotechniques 29, 52, 54. 10.2144/00291bm09.

102. Garritano, S., Gemignani, F., Voegele, C., Nguyen-Dumont, T., Le Calvez-Kelm, F., De Silva, D., Lesueur, F., Landi, S., and Tavtigian, S.V. (2009). Determining the effectiveness of High Resolution Melting analysis for SNP genotyping and mutation scanning at the TP53 locus. BMC Genet 10, 5. 10.1186/1471-2156-10-5.

103. Yamaguchi, N., Otsuna, H., Eisenberg-Bord, M., and Ramakrishnan, L. (2025). An Image Processing Tool for Automated Quantification of Bacterial Burdens in Zebrafish Larvae. Zebrafish 22, 11–14. 10.1089/zeb.2024.0170.

