## Supplementary material for "Macrophage adaptation to hypoxia in the tuberculous granuloma potentiates mycobacterium-induced mitochondrial damage and granuloma necrosis": Resources Table

| REAGENT or RESOURCE | SOURCE | IDENTIFIER |
| --- | --- | --- |
| Bacterial and virus strains | | |
| *M. marinum* M strain transformed with *pmsp12::BFP2* | Takaki et al., 2013^97^ | Derivative of ATCC # BAA-535 |

| *M. marinum* M strain transformed with *pmsp12::mWasabi* | Takaki et al., 2013^97^ | Derivative of ATCC # BAA-535 |
| --- | --- | --- |

| *M. marinum* M strain transformed with *pmsp12::tdTomato* | Takaki et al., 2013^97^ | Derivative of ATCC # BAA-535 |
| --- | --- | --- |
| *ΔESX1 M. marinum* M strain transformed with *pmsp12::tdTomato* | Pagán et al., 2015^59^ | Derivative of ATCC # BAA-535 |
| *ΔesxA M. marinum* M strain transformed with *pmsp12::tdTomato* | Osman et al., 2022^22^ | Derivative of ATCC # BAA-535 |
| *ΔesxA ::esxA^Mtb^ - M. marinum* M strain transformed with pMH406 (complementation construct containing the *M. tuberculosis esxBA* operon under control of the mycobacterial optimal promoter). | Osman et al., 2022^22^ | Derivative of ATCC # BAA-535 |
| *ΔesxA ::esxA^M83I^* - *M. marinum* M strain transformed *with* pMH406-M83I (containing M83I point mutation in esxA). | Osman et al., 2022^22^ | Derivative of ATCC # BAA-535 |
| Chemicals, peptides, and recombinant proteins | | |
| BD Difco Middlebrook 7H9 broth (dehydrated) | Fisher Scientific | Cat# DF0713-17-9 |
| Middlebrook 7H10 agar base | Sigma-Aldrich | Cat# M0303 |
| Instant Ocean Salt | ZM Systems | N/A |
| PTU (1-phenyl-2-thiourea) | Sigma-Aldrich | Cat# P7629; CAS: 103-85-5 |
| Leibovitz’s L-15 medium, no phenol red | Thermo Fisher | Cat# 21083027 |
| Tango Buffer (10x) | Thermo Fisher | Cat# BY5 |
| Phenol Red Sodium Salt | Sigma-Aldrich | Cat# P5530; CAS: 34487-61-1 |
| Pronase | Sigma-Aldrich | Cat# P5147; CAS:9036-06-0 |
| Tricaine (ethyl 3-amonobenzoate, methanesulfonic acid salt) | Fisher Scientific | Cat# 10743661; CAS: 886-86-2 |
| TopVision Low Melt Agarose | Thermo Fisher | Cat# R0801 |
| Rapamycin | Sigma-Aldrich | Cat# R0395; CAS: 53123-88-9 |

| Roxadustat | Cayman Chemical | Cat#: 15294  CAS: 808118-40-3 |
| --- | --- | --- |
| Hoechst 33342 | Tocris Biosciences | Cat# 5117  CAS: 875756-97-1 |
| MitoTracker Red CM-H2Xros | Thermo Fisher | Cat# M7513 |

| Precision Melt Supermix | BioRad | Cat# 172-5112 |
| --- | --- | --- |
| Paraformaldehyde, 16% w/v/ aqueous solution, methanol-free | Alfa Aesar | Cat# 043368.9M;  CAS: 30525-89-4 |

| KASP V4.0 2X Master Mix | LGC Biosearch | Cat# KBS-1016-002 |
| --- | --- | --- |

| Alt-R Sp Cas9 Nuclease V3 | IDT | 1081058 |
| --- | --- | --- |
| Critical commercial assays | | |
| Gibson Assembly Cloning Kit | New England BioLabs | Cat# E5510S |
| mMessage mMachine T7 Transcription Kit | Thermo Fisher | Cat# AM1344 |
| Experimental models: Organisms/strains | | |
| Zebrafish (Danio rerio): wild type AB strain | University of Cambridge | ZDB-GENO-960809-7 |
| Zebrafish: wild type TL strain | University of Cambridge | ZDB-GENO-990623-2 |
| Zebrafish: *Tg(mpeg1:Brainbow)^w201^* | Pagán et al., 2015^59^ | ZDB-ALT-150512-3 |
| Zebrafish: *Tg(mfap4:tdTomato-CAAX)^xt6^* | Walton et al., 2015^82^ | ZDB-ALT-160122-3 |
| Zebrafish: *Tg(mfap4:MTS-EGFP;myl7:RFP)^cu72^* | This work | N/A |
| Zebrafish: *Tg(mfap4:mNeonGreen-DEVD-mScarlet-I)^cu40^* | This work | N/A |
| Zebrafish: *Tg(4xtata-hre:mCherry;mlc2:eGFP)^cu73^* | This work; injected unmodified construct described in Vettori, et al. 2017^73^ | N/A |
| Zebrafish: *Tg(mfap4:DAhif1a*  *b-2A-tdTomato-CAAX)^cu74^* | This work | N/A |
| Zebrafish: Tg(*4xhre-tata:EGFP)^cu75^* | This work; injected unmodified construct described in Vettori, et al. 2017^73^ | N/A |
| Zebrafish: *Tg(-7kdrl:DsRed2*)*^pd27^* | Kikuchi, et al., 2011^108^ |  |
| Zebrafish: *vhl^sa40757^* | Wellcome Trust Sanger Institute | ZDB-ALT-160601-6912 |
| Zebrafish: *vhl^hu2117^* | Wellcome Trust Sanger Institute | ZDB-ALT-090611-1 |
| Oligonucleotides | | |
| pTol2 PhiC31LS BH New MCS (*cmlc2:RFP*) forward primer for vector fragment amplification for Gibson Assembly 5’- ATCCTCGAGCCCGGGGTT-3’ | This work | N/A |
| pTol2 PhiC31LS BH New MCS (*cmlc2:RFP*) reverse primer for vector fragment amplification for Gibson Assembly 5’-ATCCACGATCTAAAGTCATGAAGAAAGAAATCAC-3’ | This work | N/A |
| pSCAC-69 forward primer for insert fragment amplification for Gibson Assembly 5’- CATGACTTTAGATCGTGGATAAGCTTACCATGTCTGGAC-3’ | This work | N/A |
| pSCAC-69 reverse primer for insert fragment amplification for Gibson Assembly 5’- TAAACCCCGGGCTCGAGGATTTACTTGTACAGCTCGTC-3’ | This work | N/A |
| pTol2-BH:RFP-mfap4:MTS-EGFP joint sequence forward primer to validate assembled product 5’-TGAGAAGATTGCAGTAAGTT-3’ | This work | N/A |
| pTol2-BH:RFP-mfap4:MTS-EGFP joint sequence forward primer to validate assembled product  5’- CGTAGGTCAGGGTGGTCACG-3’ | This work | N/A |
| pTol2-mfap4-Lamp1-mScarleti forward primer for vector fragment amplification for Gibson Assembly  5’-ATGGTGAGCAAGGGCGAG-3’ | This work | N/A |
| pTol2-mfap4-Lamp1-mScarleti reverse primer for vector fragment amplification for Gibson Assembly  5’-CACGATCTAAAGTCATGAAGAAAG-3’ | This work | N/A |
| pT7-mNeonGreen-DEVD forward primer for insert fragment amplification for Gibson Assembly 5’-CATGACTTTAGATCGTGTAATACGACTCACTATAGGGC-3’ | This work | N/A |
| pT7-mNeonGreen-DEVD reverse primer for insert fragment amplification for Gibson Assembly 5’-GCCCTTGCTCACCATAAACTCTGATCCAGAAGTCC-3’ | This work | N/A |
| pTol2-mfap4:mNeonGreen-DEVD-mScarlet-I joint sequence forward primer to validate assembled product 5’-AAACGGAACCGAGAACTGCT-3’ | This work | N/A |
| pTol2-mfap4:mNeonGreen-DEVD-mScarlet-I joint sequence reverse primer to validate assembled product 5’-GAACTGGAGGTCACCCTTGG-3’ | This work | N/A |
| Forward primer for genotyping *vhl^hu2117^* by HRMA  5’-CGTTGAAGCTTTAGTCTAACTCGG-3’ | This work | N/A |
| Reverse primer for genotyping *vhl^hu2117^* by HRMA  5’-CGAACCCACAAAAGTTGTTATTCT-3’ | This work | N/A |
| Alt-R CRISPR Cas9.VHL.1.AA  5’-TACGTGAACATTCAGCCGTA-3’ | IDT | N/A |
| Alt-R CRISPR Cas9.VHL.1.AB  5’- GATACAGGTCAACGTTCTGT-3’ | IDT | N/A |
| Alt-R CRISPR Cas9.VHL.1.AC  5’- TCTGGATCAACTTCCTCGGA-3’ | IDT | N/A |
| Forward primer for genotyping *vhl* AA and *vhl* AC mutagenesis by HRMA  5’-AAGCCCGTCTGGATCAACTT-3’ | This work | N/A |
| Reverse primer for genotyping *vhl* AA and *vhl* AC mutagenesis by HRMA  5’- GCATAATTTCACGAACCCACA-3’ | This work | N/A |
| Forward primer for genotyping vhl AB mutagenesis by HRMA  5’- GGTCTCTGATCAGCCGGATA-3’ | This work | N/A |
| Reverse primer for genotyping vhl AB mutagenesis by HRMA  5’- CCGAGGAAGTTGATCCAGAC-3’ | This work | N/A |
| Alt-R CRISPR-Cas9 tracrRNA | IDT | 1073191 |
| Alt-R crRNA Dr.Cas9.AICDA.1.AA 5’-TCCACTATAAGAATGTGCGC-3’ | IDT | N/A |
| Forward primer for genotyping aicda AA mutagenesis by HRMA  5’-TGTGCTCATGACCCAGAAGA-3’ | This work | N/A |
| Reverse primer for genotyping aicda AA mutagenesis by HRMA  5’-TAGGTTTCGTGTCTCCCTCG-3’ | This work | N/A |
| Forward primer for *pdk1* qPCR | GCTCAGGGTGTTGTGGAATA |  |
| Reverse primer for *pdk1* qPCR | CAGAAGAGTGTGCTGGTTCA |  |
| Forward primer for *pdk2a* qPCR | ATGGTGCATAGCTGGTATATTCA |  |
| Reverse primer for *pdk2a* qPCR | GAGCAGCGACAAACTCTTCTA |  |
| Forward primer for *pdk2b* qPCR | CCAGCACACTCTGATCTTTGA |  |
| Reverse primer for *pdk2b* qPCR | GCTCTCATAGGCATCTCTGATTAC |  |
| Forward primer for *pdk3a* qPCR | CCAGCACACTCTCCTCTTTG |  |
| Reverse primer for *pdk3a* qPCR | GTAGGCATCAGTCACCACTTC |  |
| Forward primer for *pdk3b* qPCR | CTGGATTTCGGCAGGGAAA |  |
| Reverse primer for *pdk3b* qPCR | GGTGACTTCCCTCATCGTATTG |  |
| Forward primer for *pdk4* qPCR | CAATGAGAGCAACTGTGGAAAC |  |
| Reverse primer for *pdk4* qPCR | CTGCCTCTGTCAGACATCTTAAT |  |
| Forward primer for *ef1a* qPCR  5’- | CTTCTCAGGCTGACTGTGC |  |
| Forward primer for *ef1a* qPCR  5’- | CCGCTAGCATTACCCTCC |  |
| Recombinant DNA | | |
| T7-TPase | Khattak et al., 2014^101^ | RRID:Addgene_51818 |
| pSCAC-69 | Kim et al., 2008^99^ | RRID:Addgene_31241 |
| pTol2 PhiC31LS BH NewMCS (cmlc2:RFP) | Roca et al., 2019^28^ | N/A |
| pTol2 mfap4:tdTomato-CAAX | D. Tobin Laboratory; Walton et al., 2015^82^ | N/A |
| pTol2-BH:RFP-mfap4:MTS-EGFP | This work | N/A |
| pTol2-mfap4:mNeonGreen-DEVD-mScarlet-I | This work | N/A |
| Software and algorithms | | |
| NIS Elements (5.21) | Nikon | N/A |
| IMARIS for Cell Biologists (9.1) | Bitplane | N/A |
| Prism (version 10) | GraphPad | N/A |
| Image J/Fiji | https://imagej.net/software/fiji/ | N/A |
| Fluorescent Pixel Count Macro (Image J) | Takaki et al., 2013^97^ | N/A |
| Photoshop 2024 | Adobe | N/A |
| Illustrator 2024 | Adobe | N/A |
| Other | | |
| 96-well (half area) black plate with transparent bottom | Greiner Bio-One | Cat# 675090 |
| 6-well No. 1.5 coverslip, 20 mm glass diameter, uncoated plate | MatTek | Cat# P06G-1.5-20-F |
| 35 mm Dish No. 1.5 Coverslip14 mm Glass Diameter Uncoated | MatTek | Cat# P35G-0-14-C |
| Transparent WillCo-dish Glass bottom dishes, single unit packed 50 x 7 mm | WillCo Wells | Cat# GWST-5040 |
| 10 µL Microliter Syringe Model 701 N, Cemented Needle, 26s gauge, 2 in, point style 2 | Hamilton | Cat# 80366 |
